# Non-axisymmetric shapes of biological membranes from locally induced curvature

**DOI:** 10.1101/688127

**Authors:** Yannick A. D. Omar, Amaresh Sahu, Roger A. Sauer, Kranthi K. Mandadapu

## Abstract

In various biological processes such as endocytosis and caveolae formation, the cell membrane is locally deformed into curved configurations. Previous theoretical and computational studies to understand membrane morphologies resulting from locally induced curvature are often limited to axisymmetric shapes, which severely restricts the physically admissible morphologies. Under the restriction of axisymmetry, past efforts predict that the cell membrane buds at low resting tensions and stalls at a flat pit at high resting tensions. In this work, we lift the restriction of axisymmetry by employing recent theoretical and numerical advances to understand arbitrarily curved and deforming lipid bilayers. Our non-axisymmetric morphologies reveal membrane morphologies which agree well with axisymmetric studies—however only if the resting tension of the membrane is low. When the resting tension is moderate to high, we show that (i) axisymmetric invaginations are unstable; and (ii) non-axisymmetric ridge-shaped structures are energetically favorable. We further study the dynamical effects resulting from the interplay between intramembrane viscous flow and induced curvature, and find the rate at which the locally induced curvature increases is a key determinant in the formation of ridges. In particular, we show that axisymmetric buds are favored when the induced curvature is rapidly increased, while non-axisymmetric ridges are favored when the curvature is slowly increased: The rate of change of induced curvature affects the intramembrane viscous flow of lipids, which can impede the membrane’s ability to transition into ridges. We conclude that the appearance of non-axisymmetric ridges indicates that axisymmetry cannot be generally assumed when understanding processes involving locally induced curvature. Our results hold potentially relevant implications for biological processes such as endocytosis, and physical phenomena like phase separation in lipid bilayers.

## I. INTRODUCTION

The cell and its organelles are marked by a variety of strongly curved and dynamic boundaries where local curvature induction is vital. For instance, the cell membrane forms spherical vesicles as an important means of trafficking [1], and the endoplasmic reticulum maintains but also dynamically remodels networks of tubules [2, 3]. Endocytosis is another prominent biological process where curvature is locally induced. During endocytosis, proteins bend the cell membrane through different mechanisms such as scaffolding and protein insertion [4–7]. Other processes that can induce spatially varying curvatures are, for instance, charge deposition on one of the lipid monolayers [8], spatial variation of the lipid composition through phase separation [9], as well as the formation of block liposomes [10]. The examples given above show the significance of locally induced curvature in synthetic and biological systems.

While local curvature induction in lipid membranes is known to play an important role in many biological systems, the physics underlying such phenomena is not well-understood. Theoretical and numerical studies are required to better understand such processes—however, the time scale for phenomena involving local curvature induction is often on the order of seconds [11–13] and the corresponding deformations range over lengths of 100– 1000 nm [5, 11, 14]. Such length and time scales cannot be resolved using molecular simulation methods, and hence a continuum approach is often employed to understand membrane-mediated processes involving locally induced curvature.

Using a continuum approach, many studies have successfully modeled shape changes in lipid membranes. The continuum model commonly used for lipid bilayers is developed in the seminal contributions by Canham [15], Helfrich [16], and Evans [17], and can be considered an extension of Naghdi’s work on shell theory [18]. Since these pioneering developments, the model and its extensions have reproduced many experimentally observed morphologies of lipid vesicles [19–22], including tubule formation from giant unilamellar vesicles [23, 24].

Many works also studied the effects of locally induced curvature on lipid membranes. Continuum models were used in a variety of contexts, including in the study of compositional asymmetry during phase separation [25–29] and protein-induced curvature [30–34]. Biological processes such as lipid droplet formation [35, 36] and endocytosis [31, 37–42] were also modeled via locally induced curvature in previous studies. However, due to the mathematical and numerical complexity of modeling lipid membrane dynamics, most of the aforementioned studies do not allow for arbitrary deformations. Instead they are often restricted to axisymmetric shapes or small deviations from fixed geometries such as planes, cylinders or spheres. Such studies do not capture arbitrary morphological changes occurring between different geometries. Moreover, many of these studies ignore the interplay between induced curvature and intramembrane viscous flow.

The present study is based on recent theoretical advances [43–46] and corresponding numerical developments employing finite element methods [47–51], all within the framework of differential geometry, which cature the coupling between elastic out-of-plane bending and non-equilibrium processes such as intramembrane fluid flow, intramembrane phase transitions, and chemical reactions on arbitrarily curved and deforming lipid bilayers. By building on these recent advances, we study membrane morphologies resulting from locally induced curvature, without any restrictions on the permissible membrane shapes. In this work, we identify a novel, non-axisymmetric mode of deformation at moderate to high resting tensions where locally induced membrane curvature leads to the formation of ridges. In contrast, axisymmetric solutions are only preserved in the case of low resting tensions. By conducting a parameter study, we further find that the formation of ridges is influenced by the magnitude of induced curvature and its rate of increase. Thus, the assumption of axisymmetry is not generally valid in studies of locally induced curvature, and our study contradicts previous studies of membrane deformations due to locally induced curvature [37–41]. Our results advance the preliminary findings of Ref. [48], where energetically favorable, non-axisymmetric deformations were first observed.

## II. THEORETICAL MEMBRANE MODEL

In this section, we briefly describe our theoretical model; however the interested reader is referred to the Supplementary Information (SI) and Ref. [46] for further details. Lipid membranes are unique materials in that they behave like a fluid in-plane yet elastically resist bending out-of-plane. Moreover, lipid bilayers are practically area-incompressible [52]. We model the lipid membrane as a single two-dimensional manifold about the membrane mid-plane.

The elastic membrane behavior is governed by the energetic penalty for bending, commonly captured with the Helfrich free energy [16], and the membrane’s areal incompressibility. The free energy per unit area is given by

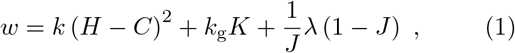

where *H* and *K* are the mean and Gaussian curvatures, respectively, *k* and *k*_g_ are the corresponding mean and Gaussian bending moduli, *J* denotes the relative change in surface area with respect to a reference configuration, and *λ* is the surface tension. In Eq. (1), the first two terms comprise the Helfrich free energy density and the last term accounts for the incompressibility constraint.

We model the effects of induced curvature with the spontaneous curvature *C*, which makes it energetically favorable for the membrane to be curved (*H*≠ 0) when-viscous ever *C* ≠ 0. However, when the membrane changes shape, lipid are required to flow in-plane. We model the in-plane flow as that of a two-dimensional Newtonian fluid, which results in additional in-plane viscous stresses (see SI). Furthermore, we note that in-plane viscous flows are coupled with the out-of-plane membrane motion [43, 44, 46], leading to an intricate relationship between surface flows, out-of-plane deformations, and surface tension gradients [44, 50]. For the length scales involved in this study, dissipation in the bulk fluid is negligible compared to the in-plane viscous dissipation [43, 53, 54]. Hence, we neglect effects of the bulk fluid surrounding the membrane. We also neglect effects from intermonolayer slip [43, 53], which are deferred to a future study.

## III. SIMULATION PROCEDURE

We employ our recent isogeometric finite element formulation [48] to simulate lipid membranes under the influence of locally induced curvature (see SI for details). To study the morphologies resulting from locally induced curvature, we consider a model system consisting of a large circular lipid bilayer patch of radius *L*, shown in gray in Fig. 1. The outer edge is subjected to a uniform surface tension *λ*_0_, which from now on will be referred to as the *resting tension*. We study local curvature induction by imposing a nonzero spontaneous curvature *C* = *C*_0_ in a chosen region in the center of the circular patch, shown in green in Fig. 1. The central patch, where *C* ≠ 0, is hereafter referred to as the *coated* area, in reference to a curvature-inducing protein coat as observed in endocytosis [6]. In all simulations, the spontaneous curvature *C*_0_ in the coated area is linearly increased from 0 to 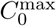 over time, at a rate *Ċ*_0_, as shown in Fig. 1. Moreover, we study the effects of varying *Ċ*_0_ on the resulting membrane morphologies.

**FIG. 1.**
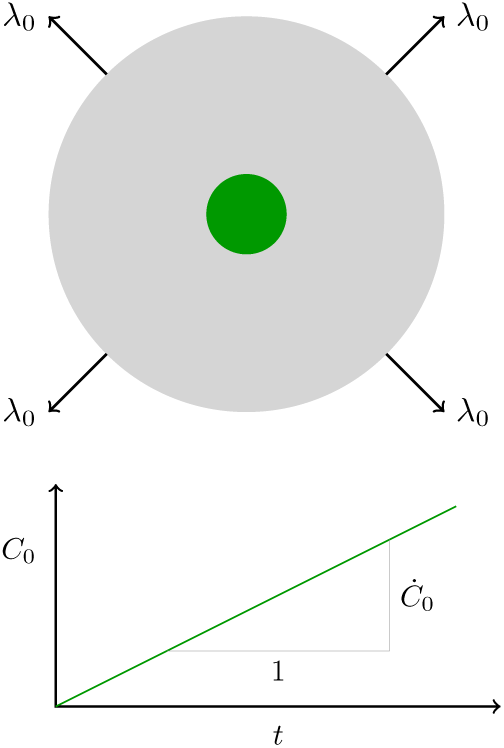
Top view of the domain used for simulations (upper inset). On the outer boundary, a boundary tension *λ*_0_ is applied to simulate the resting tension far away from the location of curvature induction. A non-zero spontaneous curvature of magnitude *C*_0_ is applied in the center of the circular geometry (shown in green), and is linearly increased over time as shown in the lower inset.

In our simulations, the coated patch is an ellipse with principal semi-axes of lengths *a* = 1.02*R*_0_ and *b* = 0.98*R*_0_, where *R*_0_ is a length that can be varied. The ellipticity breaks the symmetry of the patch, as is physically the case due to thermal fluctuations and non-circular aggregations of proteins [48]. We ensure our findings are independent of the ellipticity by considering different values of *a/b* (see SI). We assume that the coated region is convected with the surface during deformation, and neither grows nor diffuses. The latter assumption is reasonable when the timescale of diffusion of the curvature-inducing objects is much larger than the timescale of deformation of the membrane. Furthermore, curvature-inducing proteins [55, 56] and lipids [57] are known to preferentially aggregate in curved membrane regions instead of diffusing freely. In the case of clathrin-mediated endocytosis, for example, the diffusion and growth of clathrin-coated pits are negligible [5]. Moreover, during endocytosis, diffusion is further limited by the underlying actin network [58].

## IV. AXISYMMETRIC VS. NON-AXISYMMETRIC RESULTS

We now present several observations from our simulations, including both axisymmetric and non-axisymmetric results. We begin by using a numerical method to solve for axisymmetric membrane dynamics (see SI), and present results which reproduce the results of several past works [25–28, 31, 35, 37–42, 66]. We then use our general numerical framework based on finite element methods [48] and allow arbitrary membrane deformations. We find that when the resting tension is high, the non-axisymmetric shapes are significantly different from their axisymmetric counterparts, and are lower in energy. We geometrically analyze the resulting non-axisymmetric shapes and find them to be cylindrical structures. We end by using energetic arguments to justify why the non-axisymmetric structures are preferred over their axisymmetric counterparts, thus indicating the latter are unphysical in nature under certain conditions.

In all simulations presented in this section, the spontaneous curvature *C*_0_ is increased from zero to 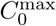 at a constant rate, for a given resting tension *λ*_0_. All results shown are instantaneous solutions, i.e. snapshots of an inherently dynamic process. We choose the rate *Ċ*_0_ to be small such that the membrane deforms slowly, the in-plane viscosity has a negligible effect on the dynamics, and the membrane generally finds its energy minimizing configuration. The material and geometric parameters chosen for our simulations are listed in Table I, which will be used hereafter unless stated otherwise.

**TABLE I.**
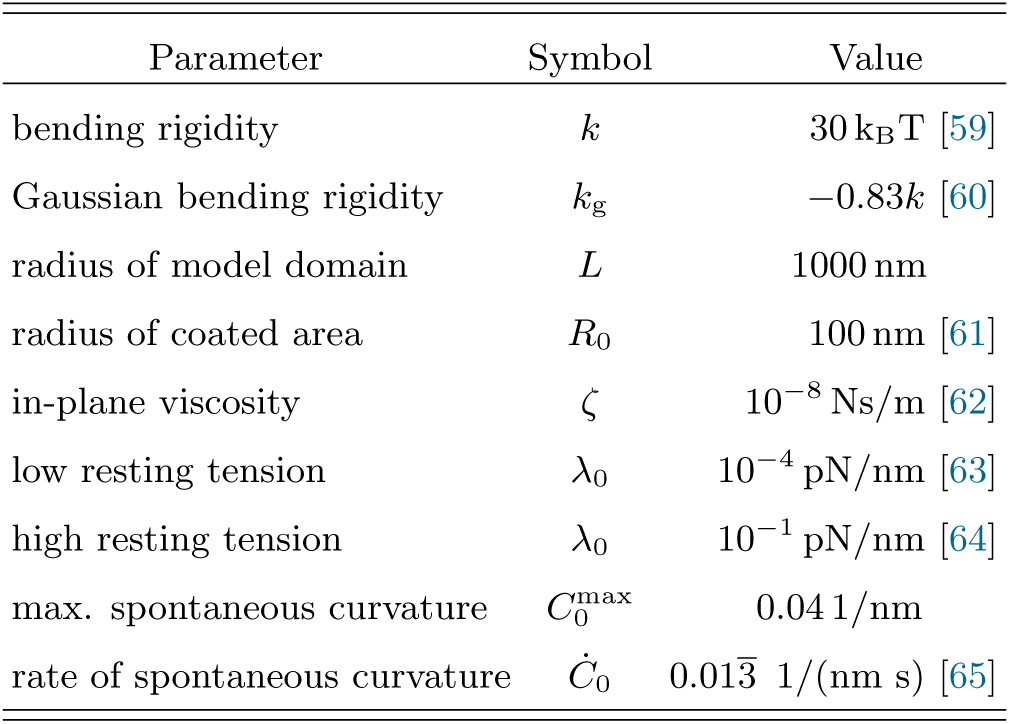
Baseline parameters used for the results of this study, unless stated otherwise. The parameters are chosen to be in biologically relevant regimes. Here, k_B_T = 4.12 pN nm, where k_B_ is the Boltzmann constant and T is the temperature.

### A. Axisymmetric Solutions

We restrict our general continuum theory [46] to axisymmetry and present a corresponding numerical method following Ref. [39] in the SI. In the axisymmetric case, the membrane’s radial, axial, and azimuthal velocities are all required to be independent of the azimuthal angle. As opposed to most axisymmetric studies of locally induced membrane curvature [25, 26, 28, 37–41, 66], we include the viscous forces arising from surface flows during membrane deformation.

Figure 2 shows our axisymmetric results at low and high resting tensions. At the low resting tension *λ*_0_ = 10^−4^ pN/nm, as the spontaneous curvature in the coated area is slowly increased, the membrane forms an invagination which deepens and eventually deforms into a bud with a constricted neck, shown in Fig. 2. On the other hand, at the high resting tension of *λ*_0_ = 10^−1^ pN/nm, the membrane forms a shallow invagination and deforms into a flat, circular pit as the spontaneous curvature is further increased. Our axisymmetric results reproduce those of earlier studies [39, 41], thus validating our numerical results.

**FIG. 2.**
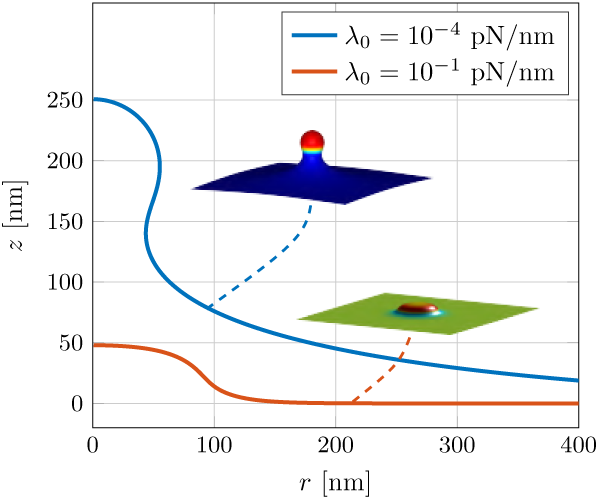
Axisymmetric shapes at different resting tensions. At a low resting tension *λ*_0_ = 10^−4^ pN/nm, the membrane forms a bud (*C*_0_ = 0.018 1/nm). When the resting tension is high, *λ*_0_ = 10^−1^ pN/nm, membrane invagination stalls—which results in a shallow, flat pit (*C*_0_ = 0.040 1/nm). The color of the three dimensional membrane configurations indicates the mean curvature *H*.

### B. Non-Axisymmetric Solutions

We next relax the constraint of axisymmetry, thus allowing general membrane deformations, using the finite element formulation developed in Ref. [48] (see SI). At the low resting tension of *λ*_0_ = 10^−4^ pN/nm, the membrane forms a shallow invagination which deepens into a bud as the spontaneous curvature is increased (Fig. 3a)—a process qualitatively similar to the axisymmetric case described above and shown in Fig. 2. In contrast, in the high resting tension case of *λ*_0_ = 10^−1^ pN/nm, the non-axisymmetric simulations differ strongly from their axisymmetric counterparts. In particular, after forming an initially shallow, axisymmetric invagination at low values of *C*_0_, the membrane deforms into a shallow horizontal ridge (Fig. 3b). The ridge is aligned along the longer principal axis of the initially elliptic patch. In what follows, we characterize the ridge structures geometrically and then provide energetic arguments why ridges are favored over the stalled, shallow pits observed in axisymmetric simulations.

**FIG. 3.**
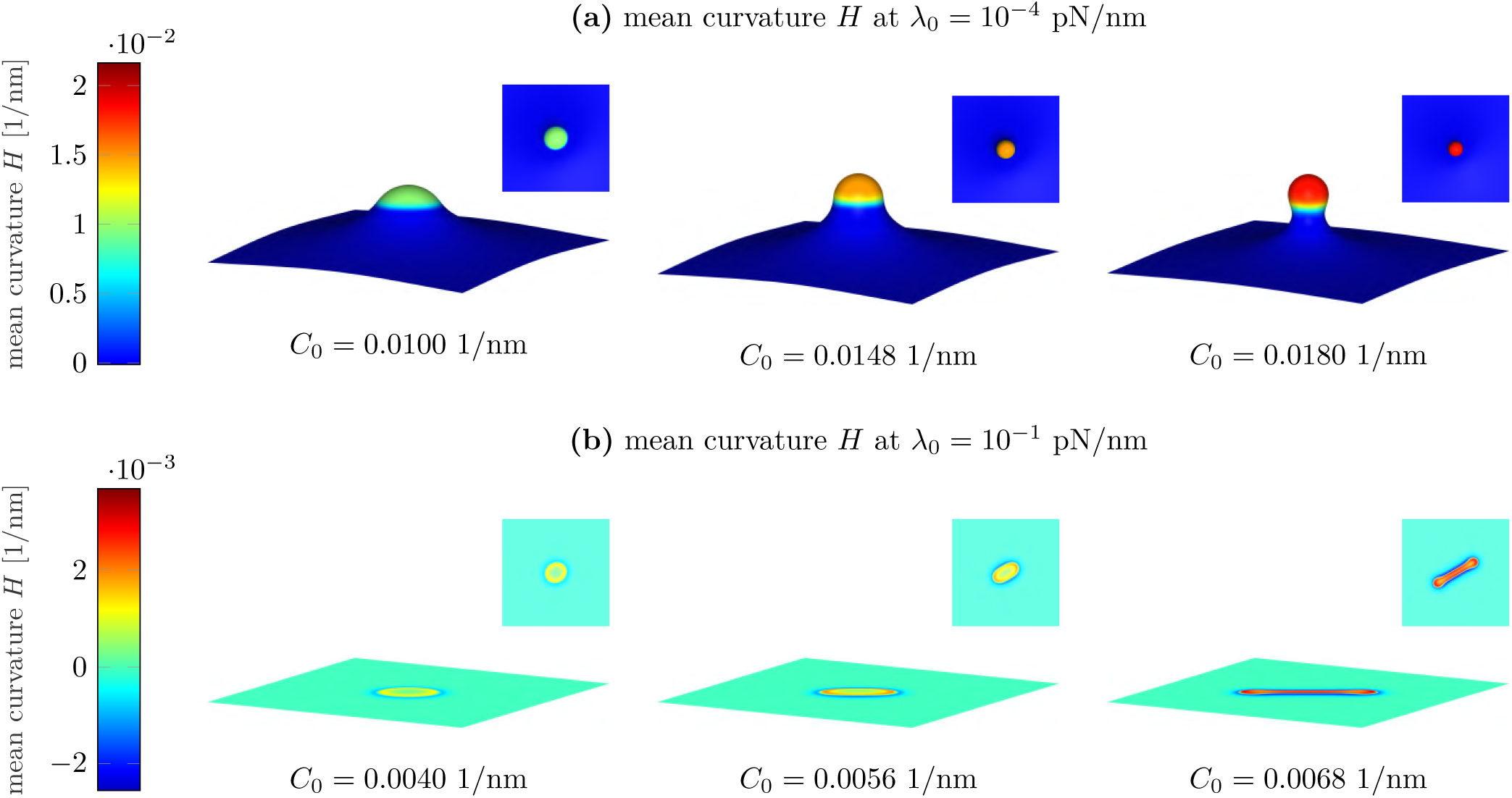
Snapshots of membrane shapes at different resting tensions *λ*_0_ and different spontaneous curvatures *C*_0_ resulting from non-axisymmetric simulations. In the low resting tension case of *λ*_0_ = 10^−4^ pN/nm, the non-axisymmetric solutions resemble the corresponding axisymmetric solutions. In the high resting tension case of *λ*_0_ = 10^−1^ pN/nm, the solution branches out into a non-axisymmetric, elongated structure, unlike its axisymmetric counterparts.

### C. Ridge Characterization

A more detailed view of the ridge geometry is shown in Fig. 4. The elongated ridge structure has a dumbbell shape and displays reflection symmetry about the principal axes of the coated region. It has a long cylindrical body and terminates in spherical caps, as shown in the zoomed-in view in Fig. 4. To further investigate the ridge geometry and compare it with the spherical buds observed at low tension, we plot the two principal curvatures *κ*_1_ and *κ*_2_ for both ridges and buds (Fig. 5). At low resting tension, *κ*_1_*/κ*_2_ is of order one in the budded region, indicating that the bud is nearly spherical.

**FIG. 4.**
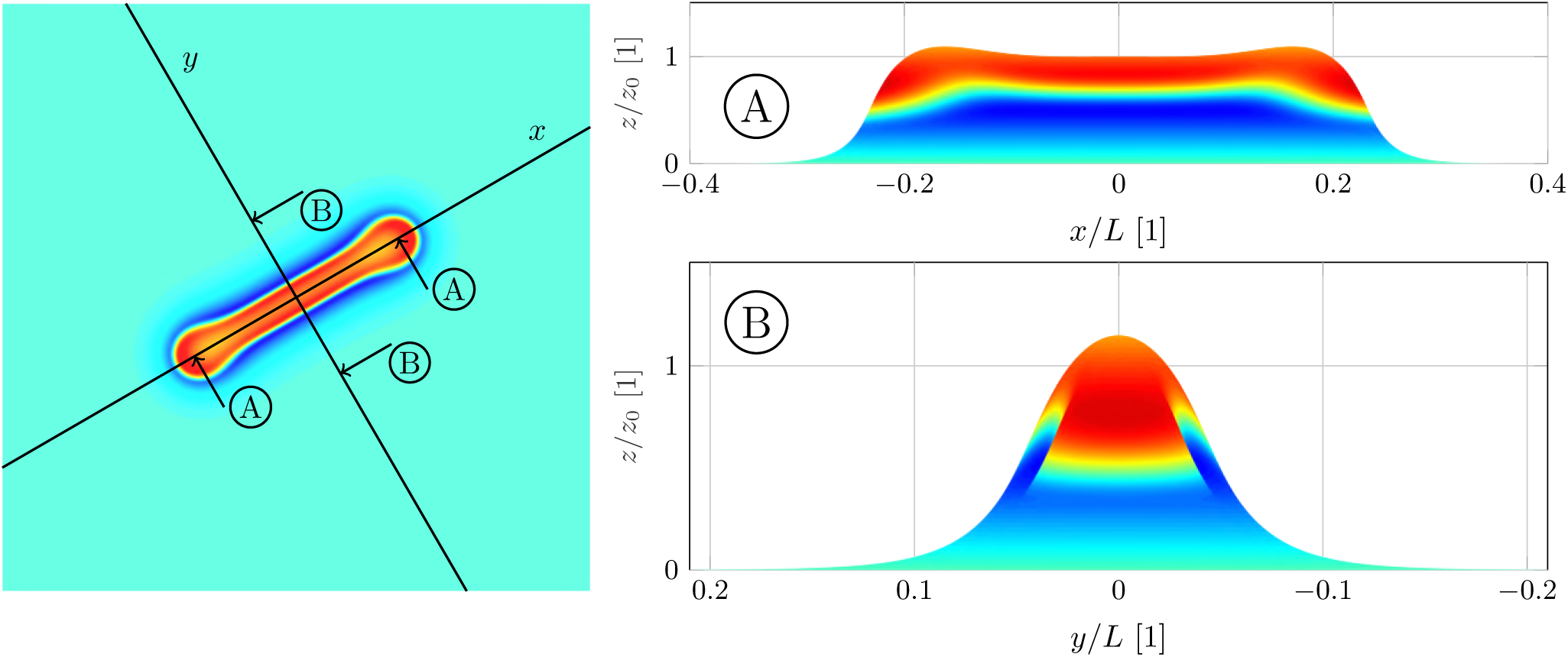
Top (left) and cross-sectional (right) views of the ridges forming at high resting tensions. The vertical axis is scaled differently from the horizontal axis, where *z*_0_:= *z*(*x* = 0*, y* = 0) = 2.6 nm, *R*_0_ = 100 nm and *L* = 1000 nm. While the geometry is not significantly curved along the longer principal axis (A), it is curved along the shorter principal axis (B), thus, resembling a cylinder. Furthermore, the spherical shape of the ends of the ridge are visible.

**FIG. 5.**
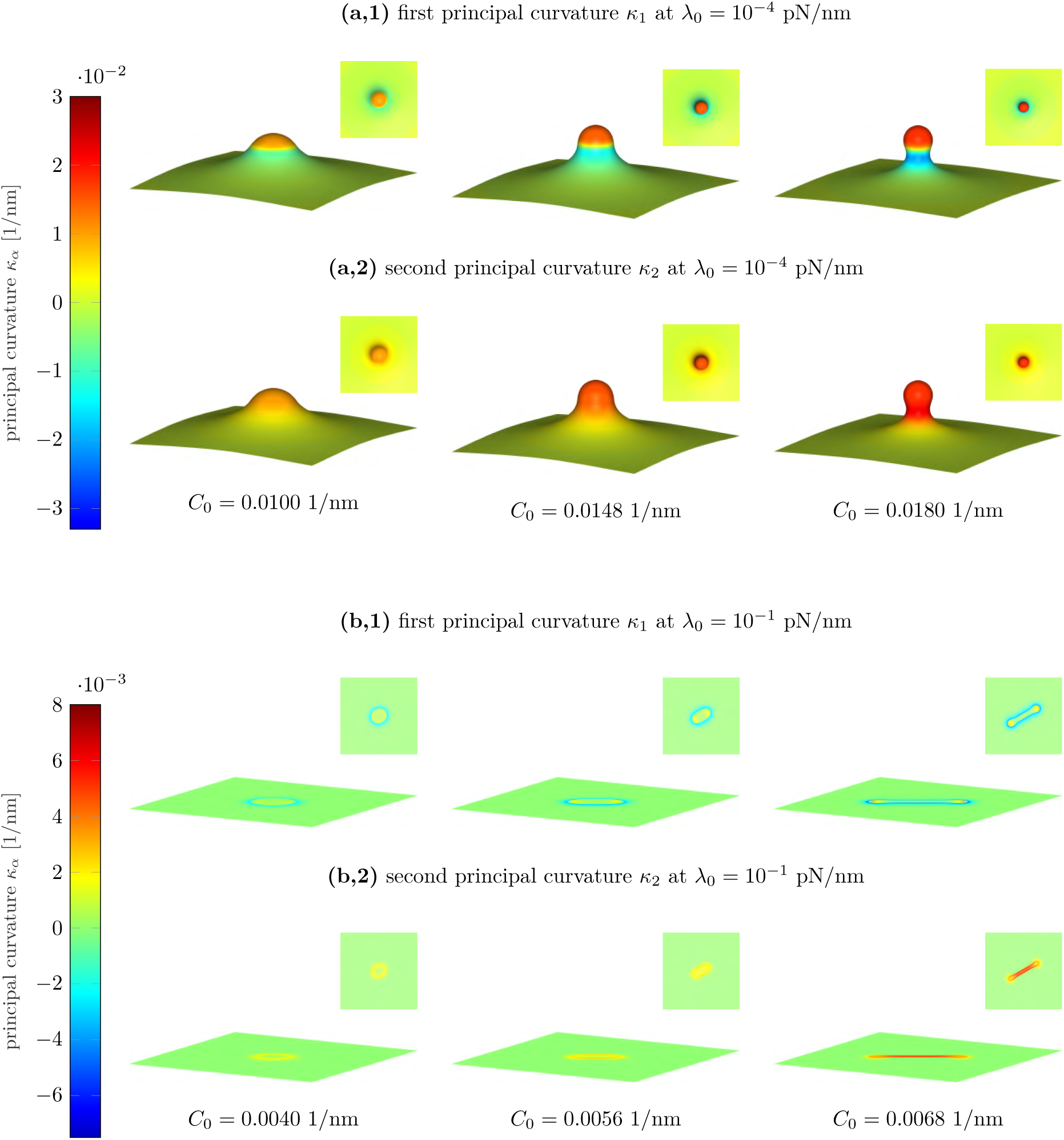
Plots of the principal curvatures *κ*_1_ and *κ*_2_ at different resting tensions *λ*_0_ resulting from the non-axisymmetric problem setup. In the low surrounding surface tension case of *λ*_0_ = 10^−4^ pN/nm, the two principal curvatures match in the region of the bud, which indicates a spherical shape. In the high resting tension case of *λ*_0_ = 10^−1^ pN/nm, the first principal curvature is one order of magnitude lower than the second principal curvature, indicating a cylindrical shape.

At high resting tension, on the other hand, the two principal curvatures are of the same order only at the beginning of the simulation, when deformations are small (Fig. 5b, left panel). As soon as the ridge develops, the first principal curvature *κ*_1_ decays to a value that is one order of magnitude lower than the second principal curvature *κ*_2_. Such a combination of principal curvatures demonstrate that the ridges are sections of cylindrical structures.

### D. Energetic Arguments: Buds vs. Ridges

To understand the difference between the axisymmetric and non-axisymmetric simulations, we consider the total elastic membrane energy

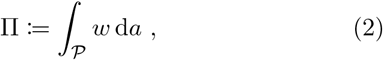

where *w* is the energy density given in Eq. (1) and the area integral is over the membrane patch 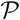. We can simplify Eq. (2) by recognizing (i) according to the Gauss– Bonnet theorem, the integral of the *k*_g_*K* term over the membrane area is a constant if the boundary remains flat, and in our case can be ignored, and (ii) the membrane is area-incompressible, such that the area stretch *J* = 1 everywhere. In this case, we can redefine our overall energy to be

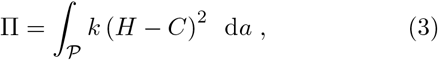

where we only need to take into account the difference between the mean and spontaneous curvatures when analyzing the membrane energetics.

We now consider the energetics of the axisymmetric and non-axisymmetric membrane shapes, which are plotted in Fig. 6. In the low resting tension case, the axisymmetric and non-axisymmetric energies are almost identical (Fig. 6), and in both cases a bud forms. However, in the high resting tension case, the axisymmetric and non-axisymmetric energies only agree at low values of *C*_0_. At higher spontaneous curvatures a ridge develops, which is lower in energy than the axisymmetric stalled pit (Fig. 6).

**FIG. 6.**
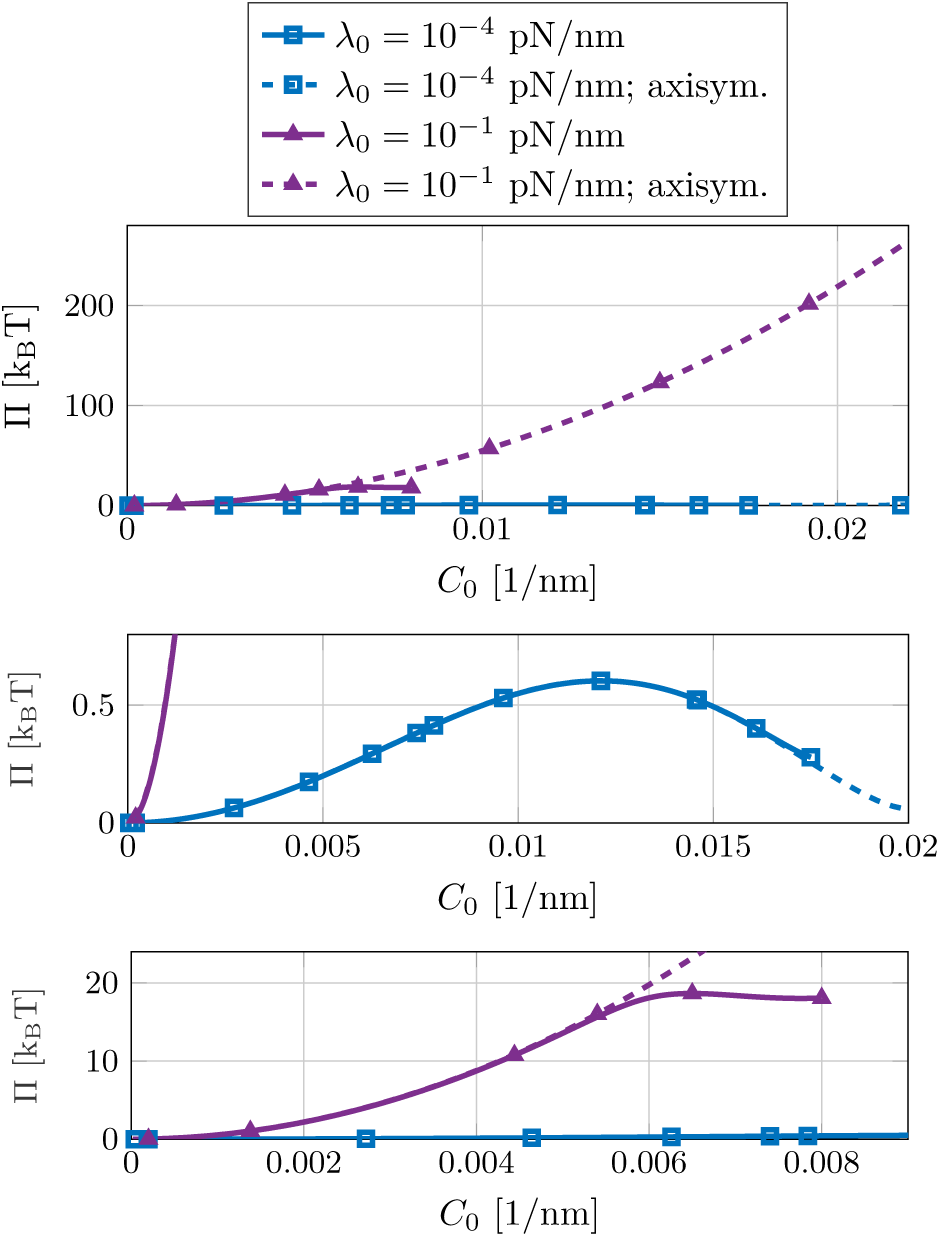
Comparison of the elastic energy Π defined in Eq. (2) for the axisymmetric and non-axisymmetric cases. The second and third subfigures show close-ups of sections of the top figure. At a low resting tensions of *λ*_0_ = 10^−4^ pN/nm, the stored energies in the two cases match closely. At a high surface tension of *λ*_0_ = 10^−1^ pN/nm, the non-axisymmetric branch deviates from the axisymmetric case and follows a path with a significantly lower energy than the axisymmetric solution. The line markers merely indicate that the plots are generated from discrete datapoints, where, for clarity, the number of line markers is much less than the number of datapoints.

At this point, we provide arguments as to why ridges are possible structures—in addition to closed buds. We begin by splitting the total membrane energy (3) into its contributions from the coated and non-coated areas, where

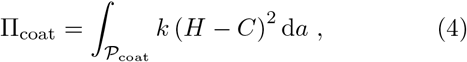

and 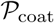 is the coated region of the membrane patch. In the coated region, we observe that *H* ≈ *C*_0_ around the resting tension and spontaneous curvature where the membrane first transitions from shallow pits to either buds or ridges. We recognize that the coat energy can be minimized when 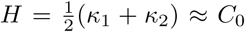 in two different ways:

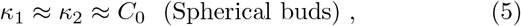

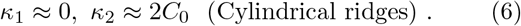

Both choices are available to non-axisymmetric simulations, while only Eq. (5) is compatible with the requirement of axisymmetry. Equation (5) leads to spherical buds and is preferred at lower tensions in both the axisymmetric and non-axisymmetric cases. On the other hand, Eq. (6) leads to cylindrical ridges and is preferred at high tensions—as seen in Fig. 5. The transition from invaginations into spherical buds or cylindrical ridges is marked by an instability (see SI). While the existence of this instability is deduced entirely from numerical experiments with the aforementioned heuristic arguments, we aim to present a detailed theoretical stability analysis in a future contribution. In particular, we seek to understand when non-axisymmetric simulations will branch into spherical buds or cylindrical ridges, as both are energy minimizing solutions (see Eqs. (5) and (6)).

Additionally, at high resting tensions, axisymmetric simulations result in shallow pits (Fig. 2), for which *H* ≈ 0 and the coat energy 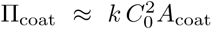 (see Eq. (4))—which is considerably larger than the energy of a ridge (see Fig. 6). Accordingly, as a much lower non-axisymmetric energy state exists, the axisymmetric results previously found at high tension [39, 41] are unstable and unphysical

## V. MORPHOLOGICAL “PHASE” DIAGRAMS

All of the results presented thus far were generated for a single coat radius *R*_0_, with the spontaneous curvature ramped up at a single rate. In this section, we explore the different morphologies accessible to lipid membranes, for a range of parameters, in both axisymmetric and non-axisymmetric settings. In particular, we study morphological “phase” diagrams, for which we consider systems (i) with different coat radii *R*_0_, (ii) over a range of resting tensions *λ*_0_, and (iii) with different rates of change of the spontaneous curvature *Ċ*_0_. We find that increasing the spontaneous curvature quickly can lead to an interplay between in-plane viscous forces and out-of-plane deformations, which prevent the membrane from reaching its lowest energy configurations—thus affecting the transitions from buds to ridges described previously. A classification of the morphologies observed in simulations is provided in Table II.

**TABLE II.** Classification of the morphologies observed in numerical simulations of locally induced curvatures. The pictographs will be used in the morphological “phase” diagrams to indicate the respective morphology. The same pictographs are used in both the axisymmetric and non-axisymmetric case for clarity.

| Number | Name | Description | Pictograph |
| --- | --- | --- | --- |
| I | pit | shallow, axisymmetric deformation |  |
| II | deep invagination | deep, axisymmetric deformation with $z(\mathbf{x} = \mathbf{0}) > R_0$ | |
| III | closed bud | deep invaginations with constricted necks |  |
| IV | ridge | non-axisymmetric, flat cylinders |  |

### A. Geometry Effects:*R*_0_ vs. *C*_0_

We first examine how different coat radii can affect membrane morphology. As shown in Fig. 7, the coat radius does not qualitatively affect the observed membrane shapes. At high resting tensions, the non-axisymmetric simulations go from shallow pits to cylindrical ridges, while the axisymmetric simulations always stall at flat, shallow pits. We note that the spontaneous curvature at which the transition occurs is almost independent of the coat radius. At low resting tensions, both axisymmetric and non-axisymmetric simulations transition from shallow pits to deep invaginations, and then to buds.

**FIG. 7.**
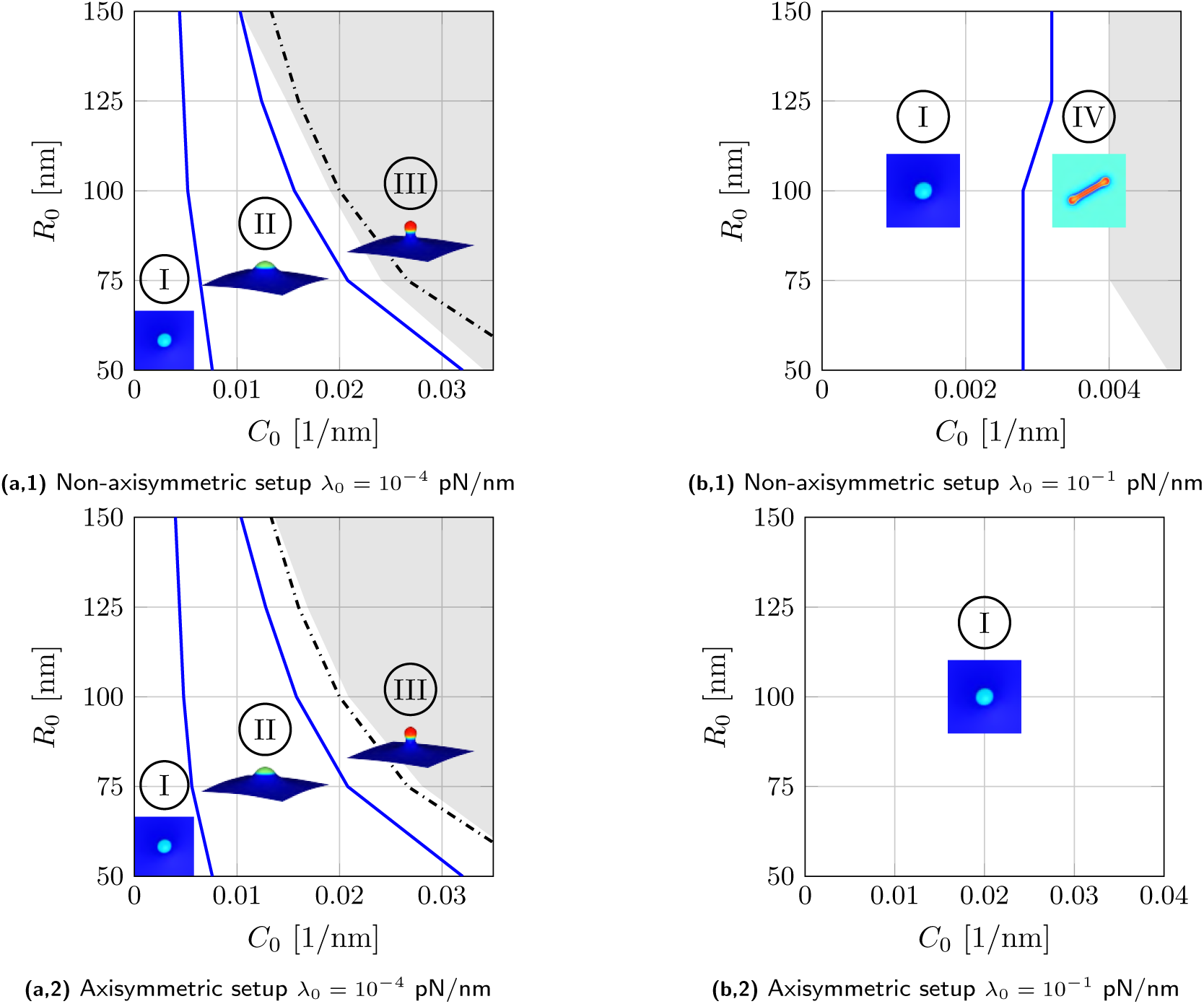
Morphological “phase” diagrams: radius of coated area *R*_0_ vs spontaneous curvature *C*_0_ at different values of the resting tension *λ*_0_ at the lowest considered rate of spontaneous curvature 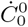. The shaded areas of the diagrams are not accessible with our current numerical framework. In the low resting tension case, an inversely proportional relation between coated area and spontaneous curvature required for closed buds is found. The dashed line shows the relationship in (7), where the proportionality was replaced by an equality. The morphological “phase” diagrams from the axisymmetric and non-axisymmetric setup agree well. In contrast to the low resting tension case, there is only a mild dependence on the size of the coated area when the resting tension is high and ridges can form. The axisymmetric simulations yield shallow pits at all spontaneous curvatures.

At low resting tension, we can also predict the onset of bud formation by considering the geometric deformation of the coated area. We reported above that for a bud, *H* ≈ *C*_0_ in the coated area (see Sec. IV D), which implies the initial coated area deforms into a spherical bud. Equating the initial and final surface areas, we find 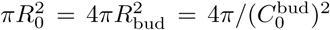, from which we approximate

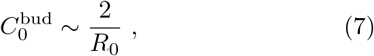

where 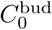 is the spontaneous curvature at which a bud is observed. Equation (7) is plotted as the dashed line in Figs. 7a,1 and 7a,2, and reasonably predicts bud formation. Accordingly, bud formation at low resting tension is a geometrical phenomenon.

### B. Resting Tension Effects: *λ*_0_ vs. *C*_0_

Thus far, we presented simulation results for two extreme cases of the resting tension: *λ*_0_ = 10^−4^ pN/nm and *λ*_0_ = 10^−1^ pN/nm. However, resting tensions in lipid membranes range from 10^−1^ pN/nm in yeast cells [39] to 3 · 10^−3^ pN/nm in blebbing cells [64, 67], and to even lower values in giant unilamellar vesicles [63]. Accordingly, we study axisymmetric and non-axisymmetric membrane morphologies over a wide range of resting tensions. Our results are captured in the morphological “phase” diagrams in Figs. 8a,1 and 8a,2, which show membrane morphologies as the spontaneous curvature is increased, for each value of the resting tension. In Figs. 8a,1 and 8a,2, the spontaneous curvature is increased slowly, such that our simulations correspond to quasi-static equilibrium configurations (Figs. 8b and 8c), discussing rate effects associated with changing *Ċ*_0_, are addressed in the subsequent section).

**FIG. 8.**
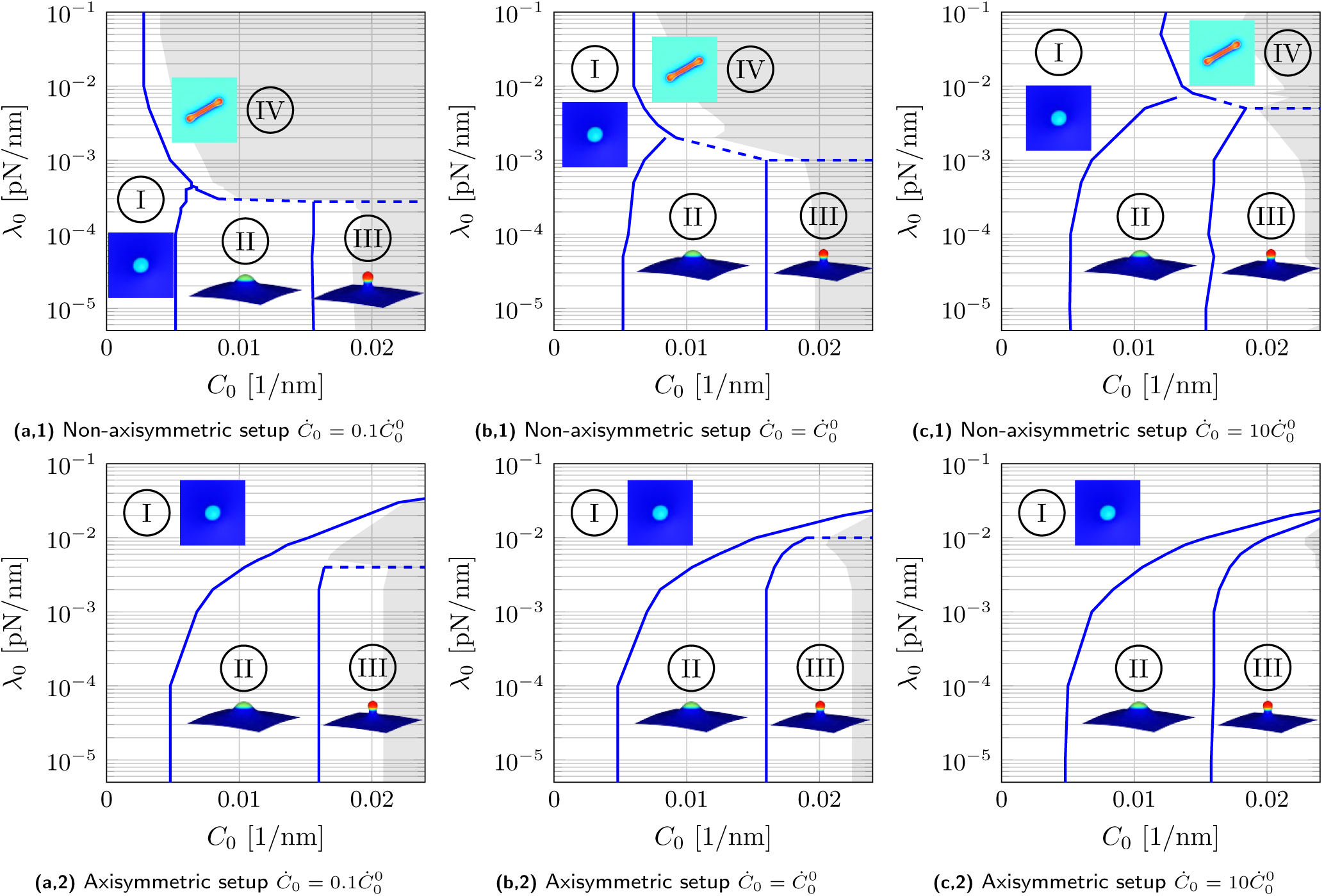
Morphological “phase” diagrams: Resting tension *λ*_0_ vs spontaneous curvature *C*_0_ at different values of the rate of spontaneous curvature *Ċ*_0_. With increased rate of spontaneous curvature, the resting tension at which ridges are observed increases. The shaded areas of the diagrams are not accessible with our current numerical framework.

The non-axisymmetric simulations shown in Fig. 8a,1 again reveal there exist two paths for morphological transitions. At resting tensions below a threshold value of *λ*_0_ ≈ 3 · 10^−4^ pN/nm, membranes transition from shallow pits to deep invaginations and then to spherical buds, with the final morphology compatible with the spherical energy minimization criterion of Eq. (5). Above this threshold, on the other hand, membranes transition from shallow pits to ridges—with the latter satisfying the cylindrical energy minimization criterion of Eq. (6). In stark contrast, the axisymmetric simulations shown in Fig. 8a,2 do not have access to the second path to form cylindrical structures; consequently, the simulations stall at shallow pits at resting tensions above 3 · 10^−4^ pN/nm.

We note that at low resting tensions, below *λ*_0_ ≈ 3 · 10^−4^ pN/nm, the axisymmetric and non-axisymmetric morphological “phase” diagrams are nearly indistinguishable, including the transitions from pits to invaginations to buds. In particular, for a given resting tension, the spontaneous curvatures at which a shallow pit becomes a deep invagination is nearly identical between the two types of simulations. The same is true for the onset of bud formation as well.

### C. Rate Effects: Varying *Ċ*_0_

In biological and artificial lipid membrane systems, the rate of curvature induction varies in different settings. For example, curvature-inducing proteins can assemble at different rates—thus inducing local curvature at different rates as well [13, 14]. In this section, we study rate effects by changing *Ċ*_0_, the rate of change of spontaneous curvature, on lipid membrane morphologies as a function of the resting tension *λ*_0_. We find that at high resting tensions, higher rates lead to additional viscous stresses in the membrane, and as a result the observed membrane morphology may not be the lowest energy configuration.

The results of changing the rate of change of spontaneous curvature, *Ċ*_0_ are presented in Fig. 8. At low resting tensions, in both the axisymmetric and non-axisymmetric cases, membrane morphologies are unaffected by changes in *Ċ*_0_: the transitions from shallow pits to deep invaginations, and then to buds, are independent of *Ċ*_0_. At high resting tensions, the axisymmetric results are largely independent of *Ċ*_0_. However, for non-axisymmetric simulations at moderate to high resting tensions, *Ċ*_0_ strongly affects membrane morphologies, as shown in Figs. 8a,1, 8b,1, and 8c,1. In particular, with increasing rates, the resting tension and spontaneous curvature at which ridges form shift toward higher magnitudes. Hence, the non-axisymmetric results increasingly resemble the axisymmetric ones when the rate of spontaneous curvature is increased (see Figs. 8c,1 and 8c,2).

To understand why the spontaneous curvature rate affects our non-axisymmetric results, we first describe the difference between axisymmetric shapes and ridges qualitatively. At high resting tensions and low rates of change of spontaneous curvature *Ċ*_0_, ridges are low energy structures, and in order to form require an in-plane shear flow of lipids. Axisymmetric shapes, on the other hand, only draw in lipids radially—thus forming shallow, flat shapes which are energetically unfavorable due to their large bending costs. We quantify the relative importance of lipid rearrangements and the forced membrane deformations by defining two relevant time scales:

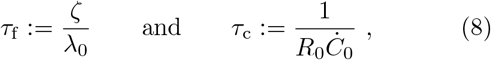

where *ζ* is the two-dimensional intramembrane viscosity, *τ*_f_ denotes the time scale associated with out-of-plane deformations and in-plane shear flows [50, 68], and *τ*_c_ is a loading time scale associated with the rate of imposed spontaneous curvature. When *τ*_f_ ≪ *τ*_c_, the lipids can quickly rearrange in-plane such that the membrane can find the lowest energy configurations—which, at high tension, are ridges. When *τ*_f_ ≫ *τ*_c_, on the other hand, lipids are unable to rearrange and access the in-plane shearing modes—and the resulting flow occurs radially in response to the changing isotropic spontaneous curvature. As the lipids cannot access the in-plane shearing modes, ridges cannot form and the membrane forms axisymmetric shapes—which are the only available option.

Our arguments on the formation of ridges and axisymmetric shapes can also explain the threshold resting tension, 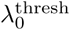, which separates ridge formation from the formation of axisymmetric shapes. This threshold is the resting tension where “phases” I, II, and IV meet (see Figs. 8a,1, 8b,1 and 8c,1). We assume the threshold value occurs when *τ*_f_ and *τ*_c_ are comparable, such that (see Eq. (8)) 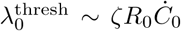. Accordingly, increasing *Ċ*_0_ by a constant factor should increase 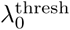 by the same factor, as *ζ* and *R*_0_ are constant. Figs. 8a,1, 8b,1, and 8c,1 show that when *Ċ*_0_ increases by a factor of 10, 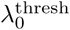 increases roughly by a factor of five. Our simple time scale argument thus predicts the correct trends for the threshold resting tension in this highly nonlinear dynamical problem. However, a detailed understanding of the effects of *Ċ*_0_ and the changes in the morphological “phase” diagrams requires a rigorous stability analysis, again involving ridge formation. We leave such an analysis to a future study.

## VI. CONCLUSIONS AND EXPERIMENTAL IMPLICATIONS

In this work, we studied lipid membrane morphologies resulting from locally-induced curvature. We found axisymmetric solutions at low resting tensions, while non-axisymmetric ridges were observed at high resting tensions. We therefore conclude that simulations which are limited to axisymmetric shapes may not provide physically meaningful results, as they cannot access lower-energy, non-axisymmetric shapes. For example, several previous studies considered the effects of locally induced spontaneous curvature as a means of studying endocytosis [39, 41]. Such works are restricted to axisymmetric shapes, and describe a snap-through instability at high tension. Our current work, however, contradicts such findings, which neglect lower energy non-axisymmetric membrane morphologies.

The non-axisymmetric lipid membrane shapes observed in the present study have implications in understanding biological processes and related phenomena. In relation to endocytosis, for example, experimental studies observe both buds at low resting tensions [12, 69] and shallow pits at high resting tensions [70, 71]. While the extended cylindrical ridges we observe at large resting tensions and spontaneous curvatures have not been explicitly reported in stalled endocytic events, there appear to be signatures of such structures in experimental studies. For example, clathrin is capable of forming cylindrical ridge-like cages in focal adhesions [72], and ridge-like polymerized structures appear to exist in clathrin-mediated endocytosis, under hypotonic conditions [71].

Ridge-like structures have also been experimentally observed on eisosomes of yeast cells [73], which generally are under high membrane tension. While eisosomes are linked to BAR proteins [73], which induce an anisotropic curvature due to their shape, our results demonstrate that even isotropic spontaneous curvatures lead to anisotropic cylindrical structures.

Additionally, a recent study found that cholera toxin subunit B (CTxB) binds to the lipid bilayer, and induces bud formation [56]. The same study suggests that bud formation is inhibited at increased resting tension, and reports ridges induced by CTxB as well [56]. However, the correlation between such ridges and the magnitude of the resting tension is not yet known.

Finally, consider morphological changes during the phase separation of biological membranes. During such processes, budding transitions were found as a result of the different spontaneous curvatures of the phase-separating components [9, 74], similar to the structures we found at low resting tension. Furthermore, there is a striking similarity between the ridge-shaped phase separated domains in lipid bilayers [9, 11], and the high resting tension ridge structures presented in our study. We speculate that such structures, as observed in Refs. [11, 75, 76], arise due to the membranes being in a high resting tension state.

## ACKNOWLEDGEMENTS

K.K.M. and Y.O. acknowledge the support of the University of California, Berkeley, and National Institutes of Health Grant No. R01-GM110066. R.A.S. acknowledges the support of the German Research Foundation (DFG) through Grant No. GSC 111 and the Aachen–California Network for Academic Exchange. A.S. is supported by the Computational Science Graduate Fellowship from the U.S Department of Energy. Y.O. was initially supported by the RWTH Scholarship for Doctoral Students and the Aachen-California Network for Academic Exchange. We are grateful to Dr. Joël Tchoufag for many stimulating discussions.

