## Supplemental Information for "Non-axisymmetric shapes of biological membranes from locally induced curvature"

(Dated: 24 July 2019)

**References** **22**

*
‡
§
†

### 1 Theory: Mechanics of Lipid Bilayers

In this section, we review the theory of the mechanics of lipid bilayers, as detailed extensively in Refs. [1,2]. We first describe the geometry and kinematics of a lipid membrane patch, then describe the membrane's constitutitive behavior, and end with the equations of motion. None of the aspects of the theoretical formulation are new, with the exception of the axisymmetric equations in Sec. 1.6, which include viscous effects.

#### 1.1 Kinematics

We begin by describing the kinematics of an arbitrarily curved and deforming surface, within the framework of differential geometry. In the following, we use the convention where subscripts and superscripts indicate covariant and contravariant components, respectively. Greek indices take values $\{1, 2\}$, and we employ the Einstein summation convention in which Greek indices repeated in a subscript and superscript are summed over.

A point $\boldsymbol{x}$ on a two-dimensional surface $\mathcal{P}$ embedded in a three-dimensional Euclidean space $\mathbb{R}^3$ is represented at time $t$ as

$$\boldsymbol{x} = \tilde{\boldsymbol{x}}(\xi^\alpha, t) = \hat{\boldsymbol{x}}(\theta^\alpha, t) = \check{\boldsymbol{x}}(\zeta^\alpha, t) \ . \tag{1}$$

Here, $\xi^\alpha$, $\theta^\alpha$, and $\zeta^\alpha$ refer to Lagrangian, in-plane Eulerian, and arbitrary Lagrangian–Eulerian descriptions (see Ref. [3]). For the sake of generality, we use the arbitrary Lagrangian–Eulerian parametrization in describing the theory. Such a parametrization allows us to represent surfaces differently based on the problem at hand. In particular, our general equations are specialized with the Lagrangian or convected coordinates, $\xi^\alpha$, for our non-axisymmetric simulations, while we use the surface-fixed coordinates, $\theta^\alpha$, in our axisymmetric simulations.

At any point $\boldsymbol{x} \in \mathcal{P}$, we define the covariant tangent vector

$$\boldsymbol{a}_\alpha := \frac{\partial \boldsymbol{x}}{\partial \zeta^\alpha} = \boldsymbol{x}_{,\alpha} \ , \tag{2}$$

where $(\bullet)_{,\alpha}$ denotes the partial derivative with respect to $\zeta^\alpha$, as well as the components of the metric tensor,

$$a_{\alpha\beta} := \boldsymbol{a}_\alpha \cdot \boldsymbol{a}_\beta \ . \tag{3}$$

The tangent vectors form a basis $\{\boldsymbol{a}_1, \boldsymbol{a}_2, \boldsymbol{n}\}$ of $\mathbb{R}^3$ (see Fig. 1) with the normal vector $\boldsymbol{n}$ given by

$$\boldsymbol{n} := \frac{\boldsymbol{a}_1 \times \boldsymbol{a}_2}{||\boldsymbol{a}_1 \times \boldsymbol{a}_2||} \ . \tag{4}$$

We define the dual basis $\{\boldsymbol{a}^1, \boldsymbol{a}^2, \boldsymbol{n}\}$, with contravariant tangent vectors $\boldsymbol{a}^\alpha$ given by

$$\boldsymbol{a}^\alpha = a^{\alpha\beta} \boldsymbol{a}_\beta \ , \tag{5}$$

which satisfy

$$\boldsymbol{a}_\alpha \cdot \boldsymbol{a}^\beta = \delta_\alpha^\beta \ , \tag{6}$$

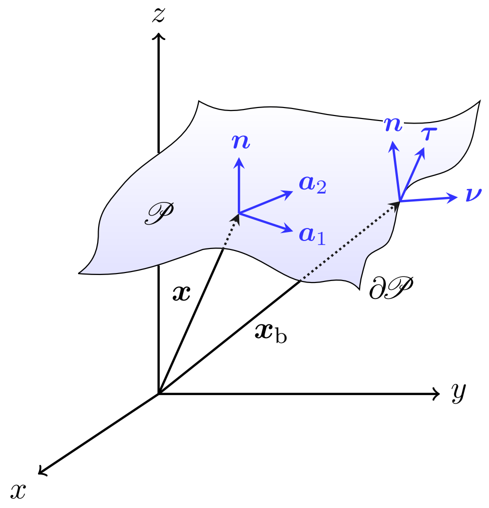

Figure 1: Schematic of a membrane patch $\mathcal{P}$. At each point $\boldsymbol{x}$ on the membrane patch $\mathcal{P}$, we define the in-plane tangent vectors $\boldsymbol{a}_1$ and $\boldsymbol{a}_2$ as well as the vector $\boldsymbol{n}$ normal to the plane spanned by $\boldsymbol{a}_\alpha$. The set $\{\boldsymbol{a}_\alpha\}$ constitutes a basis for the tangent plane, while the set $\{\boldsymbol{a}_1, \boldsymbol{a}_2, \boldsymbol{n}\}$ forms a basis of $\mathbb{R}^3$. At every point $\boldsymbol{x} \in \partial\mathcal{P}$, we define the in-plane unit tangent $\boldsymbol{\tau}$ and in-plane unit normal $\boldsymbol{\nu}$, respectively along and normal to $\partial\mathcal{P}$, which also form a basis of the tangent plane.

where the contravariant metric tensor components are defined as $a^{\alpha\beta} = (a_{\alpha\beta})^{-1}$. An arbitrary vector $\boldsymbol{h}$ can be written in the covariant and contravariant bases as

$$\boldsymbol{h} = h^\alpha \boldsymbol{a}_\alpha + h\boldsymbol{n} = h_\alpha \boldsymbol{a}^\alpha + h\boldsymbol{n} \ . \tag{7}$$

On the boundary $\partial\mathcal{P}$ of the surface $\mathcal{P}$, it is convenient to introduce a different basis $\{\boldsymbol{\nu}, \boldsymbol{\tau}, \boldsymbol{n}\}$, where $\boldsymbol{\nu}$ is the in-plane unit normal and $\boldsymbol{\tau}$ is the in-plane unit tangent at the boundary. We define

$$\boldsymbol{\tau} := \boldsymbol{a}_\alpha \frac{\partial \zeta^\alpha}{\partial \ell} \tag{8}$$

and

$$\boldsymbol{\nu} := \boldsymbol{\tau} \times \boldsymbol{n} \ , \tag{9}$$

where $\ell$ is the arclength parametrizing the patch boundary $\partial\mathcal{P}$. Next, we define the curvature tensor components

$$b_{\alpha\beta} := \boldsymbol{n} \cdot \boldsymbol{a}_{\alpha,\beta} = -\boldsymbol{n}_{,\beta} \cdot \boldsymbol{a}_\alpha \ , \tag{10}$$

and the mean and Gaussian curvatures as

$$H = \frac{1}{2}\left(\kappa_1 + \kappa_2\right) = \frac{1}{2} a^{\alpha\beta} b_{\alpha\beta} \qquad \text{and} \qquad K = \kappa_1 \kappa_2 \ = \det(b_{\alpha\beta})/\det(a_{\alpha\beta}) \ , \tag{11}$$

respectively. Figure 2 shows schematics of surface shapes with different combinations of principal curvatures, which will be useful in differentiating the morphologies identified in the main text. Figure 2a shows a spherical cap, for which the principal curvatures are equal. In contrast, the different principal axes of the ellipsoidal section in Fig. 2b yield two principal curvatures of equal sign but different magnitude. A section of a cylinder is shown in Fig. 2c, which has one vanishing

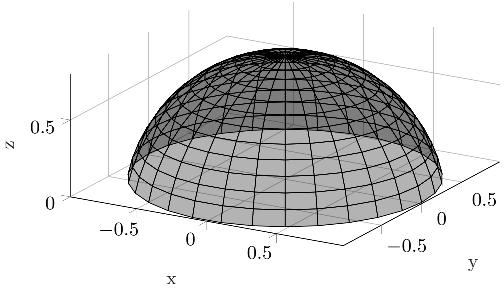

(a) Schematic of a spherical cap. Spherical sections have two equal principal curvatures.

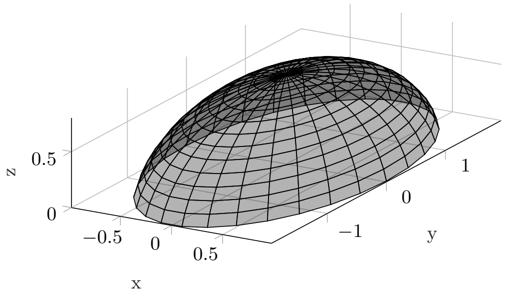

(b) Schematic of an ellipsoidal cap. Ellipsoidal sections have differing principal curvatures of the same sign.

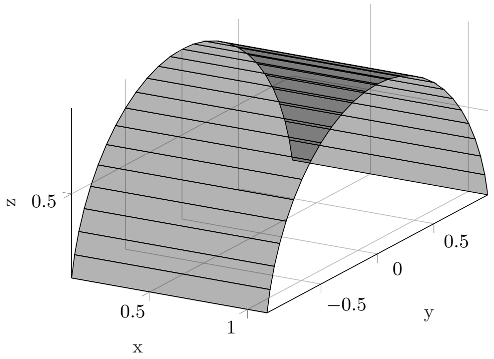

(c) Schematic of a cylindrical cap. Cylindrical sections have one vanishing and one non-zero principal curvature.

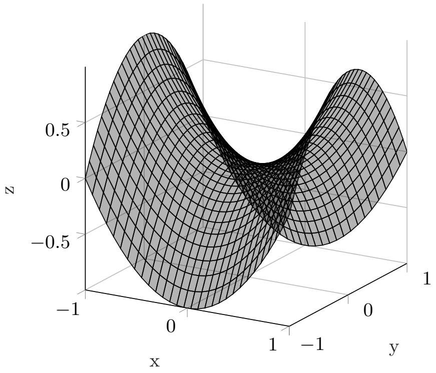

(d) Schematic of a saddle shape. Saddles are marked by two principal curvatures that differ in their sign. The saddle shown here has two equal but opposite principal curvatures such that $H = 0$ .

Figure 2: Comparison of different geometries with qualitatively different combinations of principal curvatures.

principal curvature and one non-zero principal curvature. In Fig. 2d, a saddle structure is plotted, where the two principal curvatures are equal and opposite such that the mean curvature vanishes.

The material time derivative of any quantity $(\bullet)$ is defined as the rate of change of that quantity for a given material point, written as

$$\dot{(\bullet)} := \frac{\partial(\bullet)}{\partial t}\bigg|_{\xi^\alpha} \; . \tag{12}$$

The material time derivative takes different forms in the different parametrizations defined in Eq. (1). The velocity $\boldsymbol{v} := \dot{\boldsymbol{x}}$ of any point can be written equivalently as

$$\boldsymbol{v} = \frac{\partial \tilde{\boldsymbol{x}}}{\partial t}\bigg|_{\xi^\alpha} = \frac{\partial \hat{\boldsymbol{x}}}{\partial t}\bigg|_{\theta^\alpha} + \frac{\partial \hat{\boldsymbol{x}}}{\partial \theta^\alpha}\frac{\partial \theta^\alpha}{\partial t}\bigg|_{\xi^\beta} = \frac{\partial \check{\boldsymbol{x}}}{\partial t}\bigg|_{\zeta^\alpha} + \frac{\partial \check{\boldsymbol{x}}}{\partial \zeta^\alpha}\frac{\partial \zeta^\alpha}{\partial t}\bigg|_{\xi^\beta} \; , \tag{13}$$

and can be decomposed in the $\{\boldsymbol{a}_\alpha, \boldsymbol{n}\}$ basis according to Eq. (7) as

$$\boldsymbol{v} = v^\alpha \boldsymbol{a}_\alpha + v\boldsymbol{n} \; , \tag{14}$$

where

$$v = \boldsymbol{v} \cdot \boldsymbol{n} \qquad \text{and} \qquad v^\alpha = \boldsymbol{v} \cdot \boldsymbol{a}^\alpha \ . \tag{15}$$

Moreover, the material time derivative of the covariant tangent vectors is given by

$$\dot{\boldsymbol{a}}_\alpha = \boldsymbol{v}_{,\alpha} \qquad \text{such that} \qquad \dot{a}_{\alpha\beta} = \boldsymbol{v}_{,\alpha} \cdot \boldsymbol{a}_\beta + \boldsymbol{v}_{,\beta} \cdot \boldsymbol{a}_\alpha \ . \tag{16}$$

We emphasize that the different surface parametrizations yield the same velocity $\boldsymbol{v}$ despite the functions $\tilde{\boldsymbol{x}}$, $\hat{\boldsymbol{x}}$, and $\check{\boldsymbol{x}}$ being different, and also Eq. (16) is true regardless of the surface parametrization. A more in-depth discussion of the surface parametrizations and their kinematics is provided in our previous works [1,3].

#### 1.2 Balance Laws

In the following sections, the balances of mass, linear and angular momentum will be stated in both their global and local forms. For detailed derivations, the interested reader is referred to our earlier work [1,2] and the references provided therein.

##### 1.2.1 Mass Balance

The mass balance in its global form is stated as

$$\frac{\mathrm{d}}{\mathrm{d}t} \int_{\mathcal{B}} \rho \, \mathrm{d}a = 0 \ , \tag{17}$$

where $\rho$ is the areal membrane density and $\mathcal{B}$ is an arbitrary subdomain of $\mathcal{P}$ with boundary $\partial\mathcal{B}$. Applying the Reynolds transport theorem yields, by virtue of the arbitrariness of $\mathcal{B}$,

$$\dot{\rho} + \rho(v^\alpha_{;\alpha} - 2vH) = 0 \ , \tag{18}$$

which is the local statement of mass balance—referred to as the *continuity equation*. When the material is area-incompressible, the density $\rho$ is constant and the continuity equation (18) simplifies to

$$v^\alpha_{;\alpha} - 2vH = 0 \ , \tag{19}$$

which is also known as the incompressibility constraint.

##### 1.2.2 Linear Momentum Balance

The global form of the linear momentum balance is given by

$$\frac{\mathrm{d}}{\mathrm{d}t} \int_{\mathcal{B}} \rho \boldsymbol{v} \, \mathrm{d}a = \int_{\mathcal{B}} \rho \boldsymbol{b} \, \mathrm{d}a + \int_{\partial\mathcal{B}} \boldsymbol{T} \, \mathrm{d}s \ , \tag{20}$$

with the body force per unit mass $\boldsymbol{b}$. The traction vector $\boldsymbol{T}$ at the patch boundary, due to Cauchy's triangle argument on arbitrarily curved surfaces [4], can be written as

$$\boldsymbol{T} = \boldsymbol{T}^\alpha \nu_\alpha \ , \tag{21}$$

with $\boldsymbol{T}^\alpha$ being the traction vector along $\boldsymbol{a}^\alpha$. The traction vectors can be decomposed in the $\{\boldsymbol{a}_\alpha, \boldsymbol{n}\}$ basis as

$$\boldsymbol{T}^\alpha = N^{\alpha\beta} \boldsymbol{a}_\beta + S^\alpha \boldsymbol{n} \ , \tag{22}$$

where $N^{\alpha\beta}$ and $S^{\alpha}$ are the in-plane and transverse shear components of the traction vectors, respectively. The Cauchy stress tensor can then be written as

$$\boldsymbol{\sigma} = N^{\alpha\beta}\boldsymbol{a}_{\alpha} \otimes \boldsymbol{a}_{\beta} + S^{\alpha}\boldsymbol{a}_{\alpha} \otimes \boldsymbol{n} \ , \tag{23}$$

such that $\boldsymbol{T} = \boldsymbol{\sigma}^{\mathrm{T}}\boldsymbol{\nu} = \boldsymbol{\sigma}^{\mathrm{T}}\nu_{\alpha}\boldsymbol{a}^{\alpha} = \boldsymbol{T}^{\alpha}\nu_{\alpha}$, in agreement with Eq. (21). The local form of the linear momentum balance can then be obtained as

$$\rho\dot{\boldsymbol{v}} = \rho\boldsymbol{b} + \boldsymbol{T}^{\alpha}_{;\alpha} \ . \tag{24}$$

##### 1.2.3 Angular Momentum

The global form of the angular momentum balance is written as

$$\frac{\mathrm{d}}{\mathrm{d}t}\int_{\mathcal{B}} \rho\boldsymbol{x} \times \boldsymbol{v}\,\mathrm{d}a = \int_{\mathcal{B}} \rho\boldsymbol{x} \times \boldsymbol{b}\,\mathrm{d}a + \int_{\partial\mathcal{B}} \boldsymbol{x} \times \boldsymbol{T}\,\mathrm{d}s + \int_{\partial\mathcal{B}} \boldsymbol{m}\,\mathrm{d}s \ , \tag{25}$$

where $\boldsymbol{m}$ is the moment per unit length applied on the boundary. We can express $\boldsymbol{m}$ as $\boldsymbol{m} = \boldsymbol{n}\times\boldsymbol{M}$, with $\boldsymbol{M}$ being the director traction, which can be decomposed as

$$\boldsymbol{M} = -M^{\alpha\beta}\boldsymbol{a}_{\beta} \ , \tag{26}$$

where $M^{\alpha\beta}$ are the couple-stress components (see Ref. [1]). We then find that the angular momentum balance dictates

$$1.\ S^{\alpha} = -M^{\beta\alpha}_{;\beta} \ , \tag{27}$$

$$2.\ \sigma^{\alpha\beta} := N^{\alpha\beta} - b^{\beta}_{\mu}M^{\mu\alpha} \text{ is symmetric} \ . \tag{28}$$

Note that $\sigma^{\alpha\beta}$ are the couple-free in-plane stress components.

#### 1.3 Membrane Energetics and Constitutive Behavior

The Helmholtz free energy density of a lipid membrane is denoted $\psi(a_{\alpha\beta}, b_{\alpha\beta}, T)$, where $T$ is the membrane temperature. For a general energy of this form, the membrane stresses and couple-stresses are determined via irreversible thermodynamics [1], and in the linear irreversible regime are given by

$$\sigma^{\alpha\beta} = \rho\left(\frac{\partial\psi}{\partial a_{\alpha\beta}} + \frac{\partial\psi}{\partial a_{\beta\alpha}}\right) + \pi^{\alpha\beta} \tag{29}$$

and

$$M^{\alpha\beta} = \frac{\rho}{2}\left(\frac{\partial\psi}{\partial b_{\alpha\beta}} + \frac{\partial\psi}{\partial b_{\beta\alpha}}\right) \ , \tag{30}$$

where

$$\pi^{\alpha\beta} = \zeta a^{\alpha\gamma}\dot{a}_{\gamma\delta}a^{\delta\beta} \tag{31}$$

are the viscous stresses due to in-plane incompressible flow—with $\zeta$ the in-plane dynamic viscosity.

For lipid membranes, the functional dependence of the Helmholtz free energy density can be equivalently written as

$$\psi(a_{\alpha\beta}, b_{\alpha\beta}, T) = \bar{\psi}(J, H, K, T) \ , \tag{32}$$

where $J$ denotes the relative area dilation.

For lipid membranes, the free energy consists of bending and surface tension terms—the latter arising from the incompressibility constraint. The bending energy per unit current area, called the Helfrich energy [5], is given by

$$w_h = k_{\mathrm{b}}\,(H - C)^2 + k_{\mathrm{g}} K \ , \tag{33}$$

where $C$ is the spontaneous curvature. We will discuss how the spontaneous curvature is numerically implemented in Sec. 2.3. The incompressibility of the membrane is captured by augmenting the Helmholtz free energy density with

$$w_q = \frac{1}{J}\lambda\,(J - 1) \ , \tag{34}$$

where $\lambda$ is a Lagrange Multiplier field associated with the incompressibility constraint

$$J - 1 = 0 \ . \tag{35}$$

Physically, $\lambda$ represents a surface tension. The Helmholtz free energy per unit area is then given by

$$\rho\bar{\psi} = w_h + w_q \ . \tag{36}$$

#### 1.4 General Equations of Motion

The membrane equations of motion are obtained by calculating the membrane stresses from the free energy density (36) using Eqs. (29) and (30), and then substituting the stresses into the equations of motion via Eqs. (22), (27), and (28). We present the equations of motion in component form, and to this end, the body force is split into in-plane and out-of-plane components as

$$\rho\boldsymbol{b} = p\boldsymbol{n} + b^\alpha \boldsymbol{a}_\alpha \ . \tag{37}$$

The equations of motion in the normal and in-plane directions, as well as the incompressibility constraint, are given by

$$\rho\,(v_{,t} + v^\alpha w_\alpha) = p + \pi^{\alpha\beta} b_{\alpha\beta} - 2k_{\mathrm{b}}\,(H - C)\,\big(H^2 + HC - K\big) - k_{\mathrm{b}}\Delta(H - C) + 2\lambda H \tag{38}$$

$$\rho\,\big(v^\alpha_{,t} - v w^\alpha + v^\mu w_\mu^\alpha\big) = \rho b^\alpha + \pi^{\mu\alpha}_{;\mu} - 2k_{\mathrm{b}}\,(H - C)\,C^{,\alpha} + \lambda^{,\alpha} \ , \tag{39}$$

and

$$v^\alpha_{;\alpha} - 2vH = 0 \ , \tag{40}$$

where $\Delta(\bullet) := (\bullet)_{;\alpha\beta}\, a^{\alpha\beta}$ is the surface Laplacian. Furthermore, in Eqs. (38) and (39), the inertial terms on the left-hand side use the shorthand notation

$$w_\alpha{}^\beta := v^\beta_{;\alpha} - v b^\beta_\alpha \qquad \text{and} \qquad w_\alpha := v^\lambda b_{\lambda\alpha} + v_{,\alpha} \ . \tag{41}$$

We calculate the terms in Eqs. (38) and (39) involving the viscous stresses to be

$$\pi^{\mu\alpha}_{;\mu} = 2\zeta\,\big(d^{\mu\alpha}_{;\mu} - v_{,\mu} b^{\mu\alpha} - 2vH_{,\mu} a^{\mu\alpha}\big) \tag{42}$$

and

$$\pi^{\alpha\beta} b_{\alpha\beta} = 2\zeta\,\Big[b^{\alpha\beta} d_{\alpha\beta} - 2v\,\big(2H^2 - K\big)\Big] \ , \tag{43}$$

where $d_{\alpha\beta} := \frac{1}{2}(v_{\alpha;\beta} + v_{\beta;\alpha})$ are the components of the in-plane velocity gradients.

##### 1.5 General Boundary Conditions

To solve the above equations of motion, suitable boundary conditions need to be applied. The moment and forces on the boundary in the directions $\boldsymbol{\nu}$, $\boldsymbol{\tau}$ and $\boldsymbol{n}$, respectively, are given by

$$M = k_{\mathrm{b}}\,(H - C) + k_{\mathrm{g}}\kappa_{\tau}\ , \tag{44}$$

$$f_{\nu} = k_{\mathrm{b}}\left[(H - C)^2 - (H - C)\,\kappa_{\nu}\right] - k_{\mathrm{g}}\xi^2 + \lambda + \pi^{\alpha\beta}\nu_{\alpha}\nu_{\beta}\ , \tag{45}$$

$$f_{\tau} = -\xi\left[k_{\mathrm{b}}\,(H - C) + k_{\mathrm{g}}\kappa_{\tau}\right] + \pi^{\alpha\beta}\tau_{\alpha}\nu_{\beta}\ , \tag{46}$$

$$f_{n} = -k_{\mathrm{b}}\,(H - C)_{,\nu} + k_{\mathrm{g}}\frac{\mathrm{d}\xi}{\mathrm{d}\ell}\ . \tag{47}$$

Here, $\xi$ denotes the twist of the boundary curve which is parametrized by its arclength $\ell$. At every point on the membrane boundary, we specify either the in-plane velocity components $v^{\alpha}$ or the in-plane forces $f_{\nu}$ and $f_{\tau}$. For the out-of-plane equation, we specify the membrane position and either its slope or the moment $M$. With the aforementioned boundary conditions, our equations of motion are mathematically well-posed.

##### 1.6 Axisymmetric Equations of Motion

We now present the axisymmetric equations of motion. The restriction to axisymmetry is not included in our previously published theoretical frameworks [1, 2].

Surfaces of revolution are generally parametrized by their arclength. However, when spontaneous curvature is to be prescribed locally, arclength is not a suitable parametrization to determine the coated region as the surface deforms [6, 7]. Hence, an area parametrization is chosen instead, such that the intrinsic coordinates are given by

$$\zeta^{\alpha} = \{a, \phi\}\ , \tag{48}$$

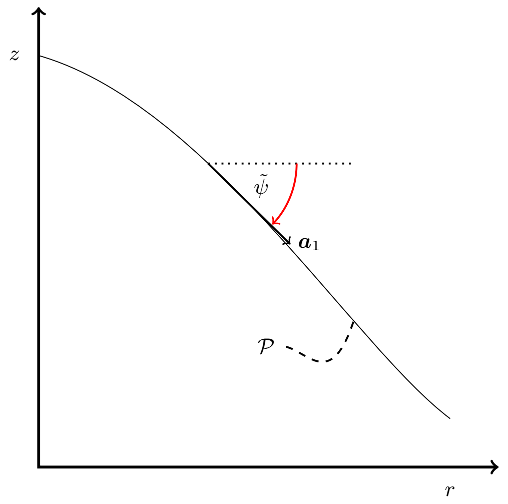

Figure 3: Schematic of the axisymmetric setup and variables. The rotationally symmetric body is represented as a line in the $r-z-$plane. The second basis vector $\boldsymbol{a}_2$ lies perpendicular to this plane.

where $a$ denotes the area parametrization—analogous to an arclength parametrization—and $\phi$ is the standard azimuthal angle.

The position of a point $\boldsymbol{x}$ on the axisymmetric surface $\mathcal{P}$ is then given by

$$\boldsymbol{x} = r\big(\zeta^1, t\big)\, \mathbf{e}_r\big(\zeta^2\big) + z\big(\zeta^1, t\big)\, \mathbf{e}_z \ , \tag{49}$$

from which the tangent and normal vectors can be computed as

$$\boldsymbol{a}_1 = r\big(\zeta^1, t\big)'\, \mathbf{e}_r\big(\zeta^2\big) + z\big(\zeta^1, t\big)'\, \mathbf{e}_z \ , \tag{50}$$

$$\boldsymbol{a}_2 = r\big(\zeta^1, t\big)\, \mathbf{e}_\phi\big(\zeta^2\big) \ , \tag{51}$$

$$\boldsymbol{n} = -\sin\tilde{\psi}\, \mathbf{e}_r + \cos\tilde{\psi}\, \mathbf{e}_z \ , \tag{52}$$

respectively, where $\tilde{\psi}$ is defined in Fig. 3 and $(\bullet)' := \partial(\bullet)/\partial\zeta^1$. The derivatives $r'\big(\zeta^1, t\big)$ and $z'\big(\zeta^1, t\big)$ are given by

$$r' = \frac{1}{2\pi r}\cos\tilde{\psi} = \tilde{J}\cos\tilde{\psi} \ , \tag{53}$$

$$z' = \frac{1}{2\pi r}\sin\tilde{\psi} = \tilde{J}\sin\tilde{\psi} \ , \tag{54}$$

with $\tilde{J} := (2\pi r)^{-1}$. The covariant components of the metric and curvature tensor, respectively, are

$$[a_{\alpha\beta}] = \begin{pmatrix} \tilde{J}^2 & 0 \\ 0 & r^2 \end{pmatrix} \qquad \text{and} \qquad [b_{\alpha\beta}] = \begin{pmatrix} \tilde{J}\tilde{\psi}' & 0 \\ 0 & r\sin\tilde{\psi} \end{pmatrix} \ . \tag{55}$$

The mean curvature and Gaussian curvature are computed from the curvature tensor to be

$$H = \frac{1}{2}\left(\frac{1}{\tilde{J}}\tilde{\psi}' + \frac{1}{r}\sin\tilde{\psi}\right) \ , \qquad \text{and} \qquad K = \frac{1}{\tilde{J}r}\tilde{\psi}'\sin\tilde{\psi} \ . \tag{56}$$

In order to obtain the viscous contributions, the velocity components need to be derived. To this end, we calculate

$$\boldsymbol{v} := \frac{\mathrm{d}\boldsymbol{x}(\zeta^\alpha, t)}{\mathrm{d}t} \ = \big(\dot{r} + r'v^a\big)\, \mathbf{e}_r + \big(\dot{z} + z'v^a\big)\, \mathbf{e}_z + rv^\phi \mathbf{e}_z \ , \tag{57}$$

where $v^a = \dot{\zeta^1}$, and $v^\phi = \dot{\zeta^2}$ . The normal velocity is given by

$$v := \boldsymbol{v} \cdot \boldsymbol{n} \ = \ \cos\tilde{\psi}\, \dot{z} - \sin\tilde{\psi}\, \dot{r} \ . \tag{58}$$

For later notational convenience, we introduce

$$\mathcal{L} := k_{\mathrm{b}}\left(\frac{r}{\tilde{J}}\,(H - C)'\right) \ , \tag{59}$$

such that

$$\mathcal{L}' = k_{\mathrm{b}} r\tilde{J}\Delta\,(H - C) \ . \tag{60}$$

The equations of motion, presented in the general case in Eqs. (38), (39), and (40), are found in the axisymmetric case to be given by

$$\begin{aligned}
&\tilde{J} r \rho \left( v_{,t} + v^a v^a \tilde{J} \tilde{\psi}' + v^a v' + v^\phi v^\phi r \sin\tilde{\psi} \right) \\
&\quad = p \tilde{J} r + 2\lambda H \tilde{J} r - 2k_{\mathrm{b}} \tilde{J} r \left( H - C \right) \left( H^2 + HC - K \right) \\
&\qquad + 2\zeta \tilde{J} r \left( 2 \frac{\tilde{\psi}'}{\tilde{J}} v H - \frac{\tilde{\psi}' r'}{\tilde{J} r} v^a + \frac{v^a}{r^2} r' \sin\tilde{\psi} - 2v \left( 2H^2 - K \right) \right) - \mathcal{L}' \ ,
\end{aligned} \tag{61}$$

$$\begin{aligned}
&\rho \left( \left( v^a \right)_{,t} - 2 v v^a \frac{\tilde{\psi}'}{\tilde{J}} - \frac{v v'}{\tilde{J}^2} + v^a (v^a)' - \frac{v^a v^a r'}{r} - v^\phi v^\phi \frac{r r'}{\tilde{J}^2} \right) \\
&\qquad = \rho b^a - 2k_{\mathrm{b}} \left( H - C \right) C' + 2\zeta \frac{\sin\tilde{\psi}}{r} \left( v' + \tilde{J} \tilde{\psi}' v^a \right) + \lambda' \ ,
\end{aligned} \tag{62}$$

$$\rho \left( \left( v^\phi \right)_{,t} - 2 v v^\phi \frac{\sin\tilde{\psi}}{r} + v^a \left( v^\phi \right)' + 2 v^a v^\phi \frac{r'}{r} \right) = \rho b^\phi + \frac{\zeta}{\tilde{J}^2} \left( \frac{3r'}{r} \left( v^\phi \right)' + \left( v^\phi \right)'' \right) \ , \tag{63}$$

and

$$0 = (v^a)' - 2vH \ . \tag{64}$$

Equations (61)–(64) are the axisymmetric shape equation, in-plane equations in the $a$- and $\phi$ direction, and the continuity equation, respectively. Equation (62) was simplified using the first derivative of the continuity equation with respect to $a$.

### 2 Numerical Methods

The following section presents the numerical approaches to solve both the axisymmetric and three-dimensional equations of motion. Our general, non-axisymmetric numerical method is based on the isogeometric, Lagrangian finite element method for arbitrarily curved and deforming surfaces developed in Ref. [8]. This method can be considered as a special case of a recently developed, more general ALE framework to describe lipid membranes [3]—which simplifies to both Lagrangian and in-plane Eulerian surface descriptions in limiting cases. Our axisymmetric equations and the corresponding simulations make use of the in-plane Eulerian description, and thus, our ALE theory provides a unified formalism to describe all of our numerics. Accordingly, due to our previous theoretical and numerical developments, we are now able to study arbitrary membrane deformations and the ensuing morphologies. We note that in all of our numerics, inertial terms are ignored.

#### 2.1 Derivation of the Axisymmetric Numerical Method

The axisymmetric equations of motion can be solved by rewriting them as a system of first order ODEs. For this purpose, we define

$$\Gamma := \left( v^\phi \right)' \ , \tag{65}$$

such that Eqs. (61)–(64) can be written as the system of equations

$$r' = \tilde{J} \cos\tilde{\psi} \ , \tag{66}$$

$$z' = \tilde{J} \sin\tilde{\psi} \ , \tag{67}$$

$$\tilde{\psi}' = \tilde{J} \left( 2H - \frac{1}{r} \sin\tilde{\psi} \right) \ , \tag{68}$$

$$H' = \frac{\mathcal{L}\tilde{J}}{k_{\mathrm{b}} r} + C' \ , \tag{69}$$

$$\begin{aligned} \mathcal{L}' = {} & p\tilde{J}r + 2\lambda H \tilde{J}r - 2k_{\mathrm{b}}\tilde{J}r \left(H - C\right)\left(H^2 + HC - K\right) + \\ & 2\zeta\tilde{J}r \left( 2\frac{\tilde{\psi}'}{\tilde{J}} vH - \frac{\tilde{\psi}' r'}{\tilde{J}r} v^a + \frac{v^a}{r^2} r' \sin\tilde{\psi} - 2v\left(2H^2 - K\right) \right) \ , \end{aligned} \tag{70}$$

$$\lambda' = -\rho b^a + 2k_{\mathrm{b}} \left(H - C\right) C' - 2\zeta \frac{\sin\tilde{\psi}}{r} \left( v' + \tilde{J}\tilde{\psi}' v^a \right) \ , \tag{71}$$

$$\left(v^\phi\right)' = \Gamma \ , \tag{72}$$

$$\Gamma' = -\frac{3r'}{r}\Gamma - \frac{\tilde{J}^2}{\zeta}\rho b^\phi \ , \tag{73}$$

$$(v^a)' = 2vH \ . \tag{74}$$

To numerically compute the time derivatives $\dot{r}$ and $\dot{z}$, an implicit finite difference scheme in time is introduced for $r(a,t)$ and $z(a,t)$:

$$\dot{\mathcal{R}}(a,t) = \frac{\mathcal{R}(a,t+\Delta t) - \mathcal{R}(a,t)}{\Delta t} + \mathcal{O}(\Delta t) \ , \tag{75}$$

$$\dot{\mathcal{R}}'(a,t) = \frac{\mathcal{R}'(a,t+\Delta t) - \mathcal{R}'(a,t)}{\Delta t} + \mathcal{O}(\Delta t) \, , \tag{76}$$

where $\mathcal{R} \in \{r, z\}$. For the latter, sufficient continuity is required such that

$$\left(\dot{\mathcal{R}}\right)' = \dot{\overline{(\mathcal{R})'}} \ , \tag{77}$$

where the horizontal bar indicates to which term the time derivative is applied.

#### 2.2 General, Non-Axisymmetric Numerical Method

The numerical method presented in the following is based on Ref. [8], to which we refer the reader for additional details. Since we employ a Lagrangian parametrization of the surface, all of our integrals over the current membrane surface are converted into integrals over the reference membrane configuration.

##### 2.2.1 Weak Forms

We begin by revisiting the mass balance given in (17) and note that we can analogously state

$$\int_{\mathcal{B}} \rho \, \mathrm{d}a = \int_{\mathcal{B}_0} \rho_0 \, \mathrm{d}a \qquad \forall t \in [0, T] \ , \tag{78}$$

where $\mathcal{B}_0$ denotes a subset of the membrane's reference configuration $\mathcal{P}_0$, and $\rho_0$ denotes the corresponding reference areal mass density. By assuming incompressibility, Eq. (78) becomes

$$J = \frac{\rho_0}{\rho} = 1 \; . \tag{79}$$

Multiplying (34) by a variation $\delta\lambda \in \mathcal{Q}$, where $\mathcal{Q}$ is the space of admissible functions for $\delta\lambda$, and integrating over the membrane surface yields

$$G_\mathrm{g} = \int_{\mathcal{P}_0} \delta\lambda\,(J-1)\,\mathrm{d}A \; . \tag{80}$$

Thus, the incompressibility constraint is satisfied if $G_\mathrm{g} = 0$ for any admissible variation $\delta\lambda$. Next, the linear momentum balance in (24) is contracted with a suitable variation $\delta\boldsymbol{x} \in \mathcal{V}$, where $\mathcal{V}$ is another admissible space. The resulting weak form is then split into the three contributions

$$G_\mathrm{inertia} = \int_{\mathcal{P}} \delta\boldsymbol{x}\cdot\rho\dot{\boldsymbol{v}}\,\mathrm{d}a \; , \tag{81}$$

$$G_\mathrm{int} = \int_{\mathcal{P}} \frac{1}{2}\delta a_{\alpha\beta}\sigma^{\alpha\beta}\,\mathrm{d}a + \int_{\mathcal{P}} \delta b_{\alpha\beta}M^{\alpha\beta}\,\mathrm{d}a \; , \tag{82}$$

$$G_\mathrm{ext} = \int_{\mathcal{P}} \delta\boldsymbol{x}\cdot\rho\boldsymbol{b}\,\mathrm{d}a + \int_{\partial\mathcal{P}} \delta\boldsymbol{x}\cdot\boldsymbol{T}\,\mathrm{d}s + \int_{\partial\mathcal{P}} \delta\boldsymbol{n}\cdot\boldsymbol{M}\,\mathrm{d}s \; . \tag{83}$$

Since $G_\mathrm{g} \equiv 0$, the weak formulation of the problem can be written as

$$G_\mathrm{g} + G_\mathrm{inertia} + G_\mathrm{int} - G_\mathrm{ext} = 0 \; . \tag{84}$$

###### 2.2.2 Finite Element Discretization

We introduce a non-overlapping discretization of the reference domain $\mathcal{P}_0$ as

$$\mathcal{P}_0 = \bigcup_{e=1}^{n_\mathrm{el}} \Omega_0^e \; , \tag{85}$$

where $\Omega_0^e$ refers to a single element $e$. A position $\boldsymbol{x} \in \mathcal{P}$ and the Lagrange multiplier as well as their variations are then approximated as

$$\boldsymbol{x}^h = \mathbf{N}^{\boldsymbol{x}}\mathbf{x} \; , \tag{86}$$

$$\lambda^h = \mathbf{N}^{\lambda}\boldsymbol{\lambda} \; , \tag{87}$$

$$\delta\boldsymbol{x}^h = \mathbf{N}^{\boldsymbol{x}}\delta\mathbf{x} \; , \tag{88}$$

$$\delta\lambda^h = \mathbf{N}^{\lambda}\delta\boldsymbol{\lambda} \; . \tag{89}$$

In Eqs. (86)–(89), $\mathbf{x}$ and $\boldsymbol{\lambda}$ are the discretized position and surface tension degree of freedom vectors and $\mathbf{N}^{\boldsymbol{x}}$ and $\mathbf{N}^{\lambda}$ are the corresponding finite element method shape functions written in array form.

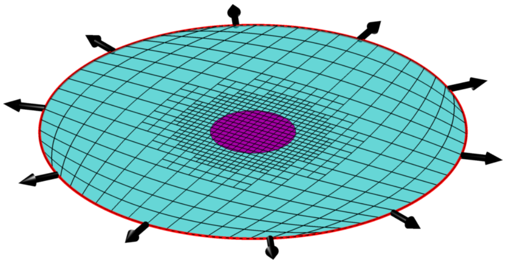

Figure 4: Domain used for simulations. The circular geometry is discretized by finite elements. On the outer boundary, a boundary tension is applied to simulate the resting tension far away from the region of non-zero spontaneous curvature. The outer boundary is constrained to only displace within the plane defined by $\mathcal{P}_0$. A non-zero spontaneous curvature is applied in the center of the circular geometry and is linearly increased over time.

###### 2.2.3 Discretized Weak Form and Linearization

In the following, it is assumed that inertial terms are negligible, such that $G_{\text{inertia}} \approx 0$. In this subsection, we omit the superscript $h$ of the interpolated quantities for clarity. Inserting the interpolations (86)–(89) into the weak forms in (84) then yields

$$\delta\mathbf{x} \cdot (\mathbf{f}_{\text{int}} - \mathbf{f}_{\text{ext}}) + \delta\boldsymbol{\lambda} \cdot \mathbf{f}_{\text{q}} = 0 \ . \tag{90}$$

By defining the residual vector

$$\mathbf{R}(\mathbf{x}, \boldsymbol{\lambda}) := \Big[(\mathbf{f}_{\text{int}} - \mathbf{f}_{\text{ext}})^{\text{T}} \ , \ \mathbf{f}_{\text{q}}^{\text{T}}\Big]^{\text{T}} \ , \tag{91}$$

the solution is obtained by solving

$$\mathbf{R}\big(\mathbf{x}, \boldsymbol{\lambda}\big) = \mathbf{0} \ . \tag{92}$$

Equation (92) is nonlinear and hence solved by using the classical Newton-Raphson scheme

$$\mathbf{R}\big(\mathbf{x}^{\text{n}+1}, \boldsymbol{\lambda}^{\text{n}+1}\big) \approx \mathbf{R}(\mathbf{x}^{\text{n}}, \boldsymbol{\lambda}^{\text{n}}) + \nabla\mathbf{R}\big|_{\mathbf{x}^{\text{n}}, \boldsymbol{\lambda}^{\text{n}}} \Delta\mathbf{u} \ , \tag{93}$$

where $\Delta\mathbf{u}$ denotes the increment in $\mathbf{x}$ and $\boldsymbol{\lambda}$.

###### 2.2.4 Meshing

The elements employed here are based on isogeometric finite elements such that the interpolation functions are Non-Uniform Rational B-Splines (NURBS). In an attempt to satisfy the LBB-condition associated with the incompressibility constraint [9–11], we interpolate $\boldsymbol{x}$ with third order polynomials and $\lambda$ with first order polynomials. Despite this, our scheme is not LBB-stable. However we supress the LBB-instability by fixing the resting tension everywhere on the boundary—which, while not sufficiently general, is suitable for our calculations [8]. In particular, we did not observe any surface tension oscillations, which are characteristic for LBB-unstable schemes. However, as shown in Ref. [3], the LBB-instability can be properly eliminated within our computational framework, as will be done in future studies.

We mesh the membrane surface with the recently introduced Locally-Refined NURBS (LR-NURBS) [12]. Elements that lie within a circular region of interest, where the spontaneous curvature is imposed, are refined multiple times with a transition region towards coarser elements in the peripheral region (see Fig. 4).

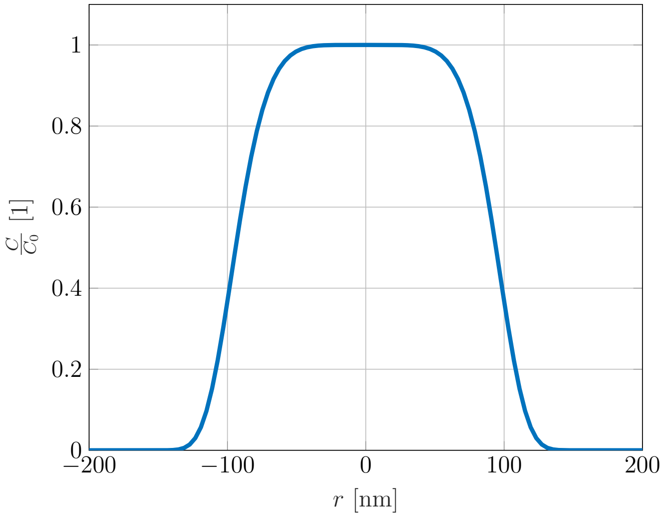

Figure 5: The spontaneous curvature distribution is shown for $R_0 = 100$ nm. It is chosen similarly to Ref. [7].

##### 2.3 Application of Spontaneous Curvature

We enforce a non-zero spontaneous curvature only in a chosen patch given by the distribution of spontaneous curvature

$$C(\tilde{X}_1, \tilde{X}_2) = C_0 \mathrm{e}^{-\left(\tilde{X}_1^2 + \tilde{X}_2^2\right)^{\frac{\beta}{2}}} , \tag{94}$$

with

$$\tilde{X}_1 = \frac{\cos\Phi\left(X_1 - X_1^0\right) + \sin\Phi\left(X_2 - X_2^0\right)}{a} , \tag{95}$$

$$\tilde{X}_2 = \frac{-\sin\Phi\left(X_1 - X_1^0\right) + \cos\Phi\left(X_2 - X_2^0\right)}{b} , \tag{96}$$

where the superscript 0 indicates the coordinates of the center of the patch, $a$ and $b$ are the length of the principal semi-axes, and the angle $\Phi$ describes the rotation of the elliptic patch. We note that the spontaneous curvature is defined by the coordinates of a material point in the reference configuration. The spontaneous curvature distribution with principal semi-axes length $a = b = R_0$ is shown in Fig. 5, for the case of $R_0 = 100$ nm. In all of our simulations in the main text, we choose the two principal axes to be of slightly different lengths: $a = R_0(1 + \delta)$ and $b = R_0(1 - \delta)$, where $\delta = 0.02$ (see Sec. 3.2.1).

#### 3 Convergence Study

In the following, we examine the convergence of the axisymmetric and non-axisymmetric simulations with respect to the time step and the mesh size. All convergence results are shown in both the low and high resting tension cases. Convergence is evaluated with the total Helfrich elastic energy (c.f. Eq. (33))

$$\Pi := \int_{\mathcal{P}} k_{\mathrm{b}} \left(H - C\right)^2 \mathrm{d}a , \tag{97}$$

as the surface tension does not contribute to the energy since $J = 1$ everywhere. The Gaussian curvature term can be neglected according to the Gauss-Bonnet theorem as is discussed in the main text. Note that in all results presented here, parameters are chosen as specified in Table I of the main text.

##### 3.1 Axisymmetric Convergence

The convergence of our axisymmetric simulations with respect to the number of grid points is shown in Fig. 6. In all cases, the spacing between grid points decreases as we approach the axis of symmetry. We find that in both the low and high resting tension cases, simulation results are largely independent of the number of grid points, $\bar{m}$. Hence, even the coarsest mesh is a reasonable choice for our simulations.

We also consider the convergence of the axisymmetric simulations with respect to the timestep size $\Delta t$, as shown in Fig. 7. Such calculations were repeated for different rates of change of spontaneous curvature, but the results are omitted here for brevity.

##### 3.2 Non-Axisymmetric Convergence

In what follows, we investigate $h$-refinement of the solutions from our general finite element framework. We again study the convergence at two different resting tensions: the low resting tension of $\lambda_0 = 10^{-4}$ pN/nm where buds form, and the high resting tension of $\lambda_0 = 10^{-1}$ pN/nm where ridges form. Results are plotted with respect to the number of local mesh refinements, $n_{\mathrm{ref}}$ in the center of the geometry. Details of our meshing procedure can be found in Ref. [12], and a mesh with $n_{\mathrm{ref}} = 3$ is shown in Fig. 4. We note that the mesh is refined prior to the simulation, and is thus not adaptive. We also studied the convergence of the outer mesh elements, with the results omitted here for brevity.

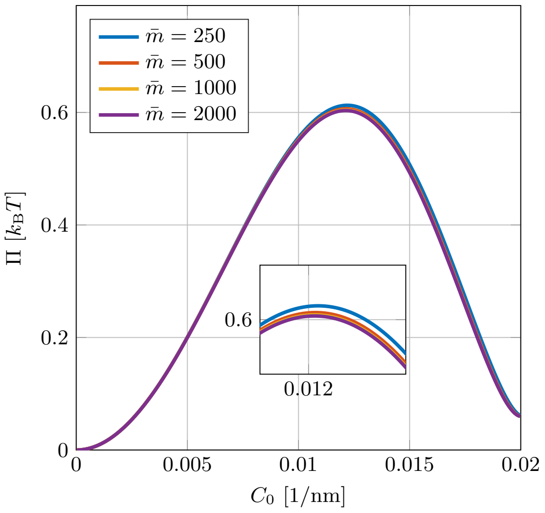

(a) at $\lambda_0 = 10^{-4}$ nm.

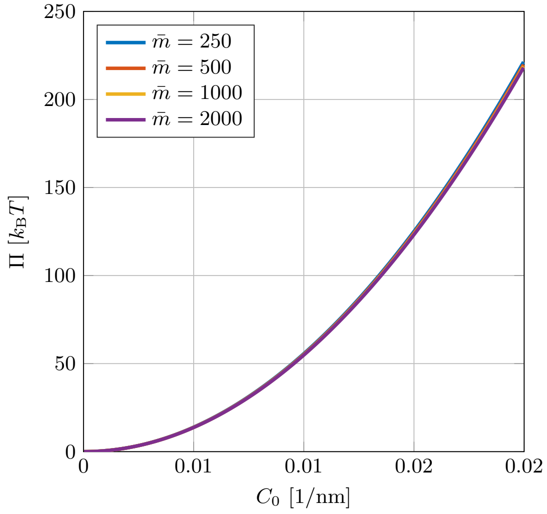

(b) at $\lambda_0 = 10^{-1}$ nm.

Figure 6: Stored energy depending on the number of grid points at different resting tensions in the axisymmetric setup.

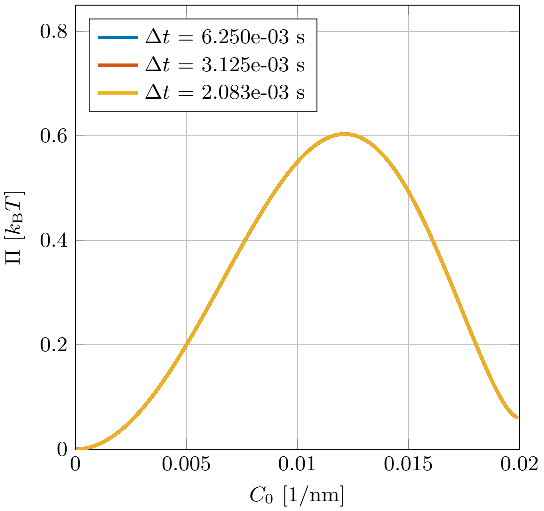

(a) at $\lambda_0 = 10^{-4}$ nm.

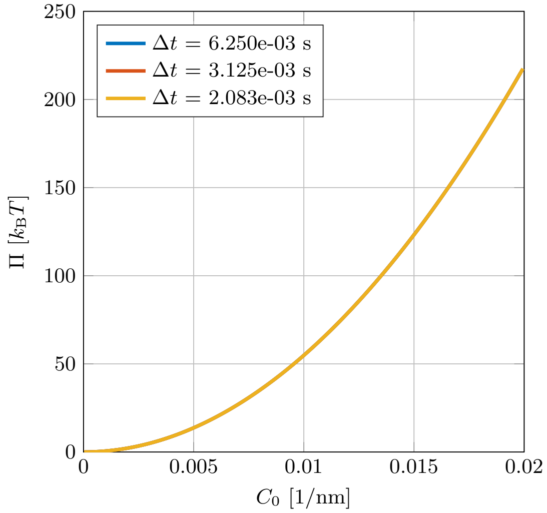

(b) at $\lambda_0 = 10^{-1}$ nm.

Figure 7: Stored energy depending on the number of timesteps at different resting tensions in the axisymmetric setup.

**Mesh Refinement: Low Resting Tension.** Convergence results for the low resting tension case are shown in Fig. 8a. With $n_{\text{ref}} = 3$, the solution appears to be converged over a large range of values for $C_0$. However, at $C_0 \approx 0.018$ 1/nm, the solution branches into a higher energy path—corresponding to a twisted bud, as discussed further in Sec. 3.2.2. In our study, no results beyond this divergence point are reported.

**Mesh Refinement: High Resting Tension.** Convergence results from the ridge-forming, high resting tension case are shown in Fig. 8b. The solution is converged for all refinement steps $n_{\text{ref}}$ shown. However, at $C_0 \approx 0.008$ 1/nm the spherical cap at each end of the ridge splits to form another ridge, as shown in Fig. 9. The formation of these branched ridges, however, is mesh-dependent and care must be applied to understand membrane morphologies involving such ridges. Furthermore, as will be discussed in detail in Sec. 3.2.1, when spontaneous curvature is applied on a strictly circular patch, the non-axisymmetric simulations result in branched ridge structures. However, these branched ridge structures depend on the choice of the mesh—as is the case for branched ridges forming from elliptic patches. We leave the analysis of instabilities involving branched ridges to a future study.

**Mesh Refinement: Refinement Region.** The size of the center region in which mesh elements are refined is defined by the radius of refinement $r_{\text{ref}}$. In both the low and high resting tension cases, we found converged results for all refinement regions considered (Fig. 10).

**Time Step Refinement.** Convergence with respect to the timestep size is shown in Fig. 11, and indicates that results are converged with respect to changing $\Delta t$ for the entire range of tensions considered. This result was reproduced for different rates of change of the spontaneous curvature, however, the results are omitted here for brevity.

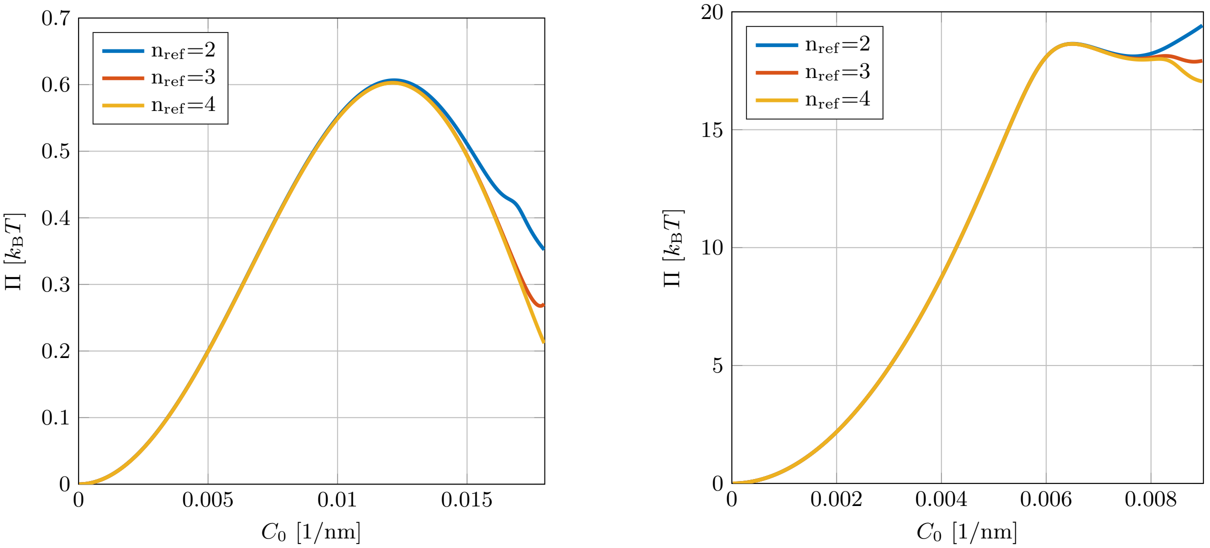

(a) at $\lambda_0 = 10^{-4}$ nm.

(b) at $\lambda_0 = 10^{-1}$ nm.

Figure 8: Stored energy depending on the number of refinement steps in the center of the computational domain at different resting tensions in the non-axisymmetric setup.

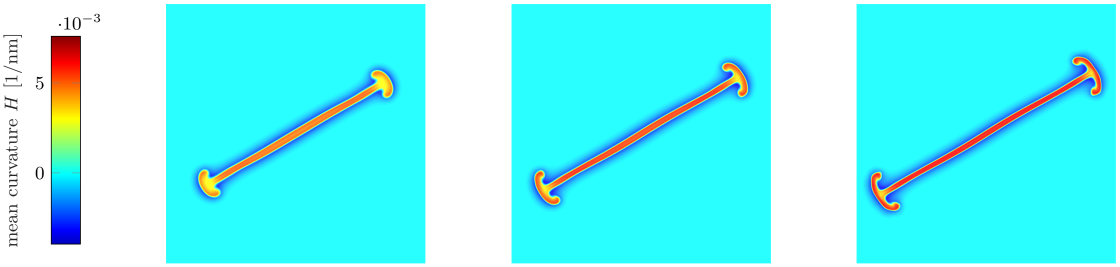

$C_0 = 0.0084$ 1/nm

$C_0 = 0.0088$ 1/nm

$C_0 = 0.0092$ 1/nm

Figure 9: When the spontaneous curvature is increased further after the ridge has formed, the spherical caps of the ridge undergo another branching. This instability is of the same nature as the initial ridge formation.

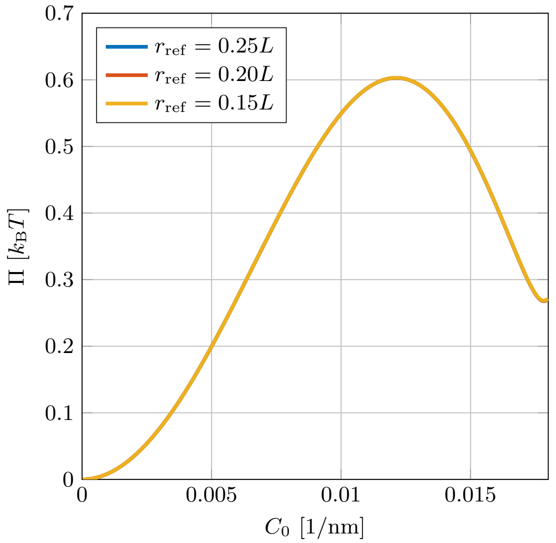

(a) at $\lambda_0 = 10^{-4}$ nm.

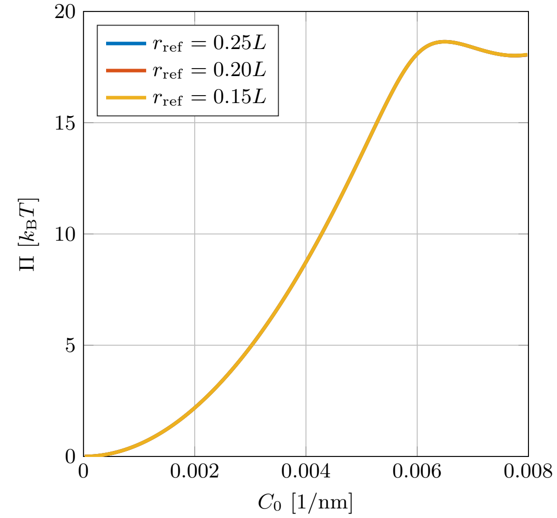

(b) at $\lambda_0 = 10^{-1}$ nm.

Figure 10: Stored energy depending on the size of the refined area in the center of the computational domain at different resting tensions in the non-axisymmetric setup.

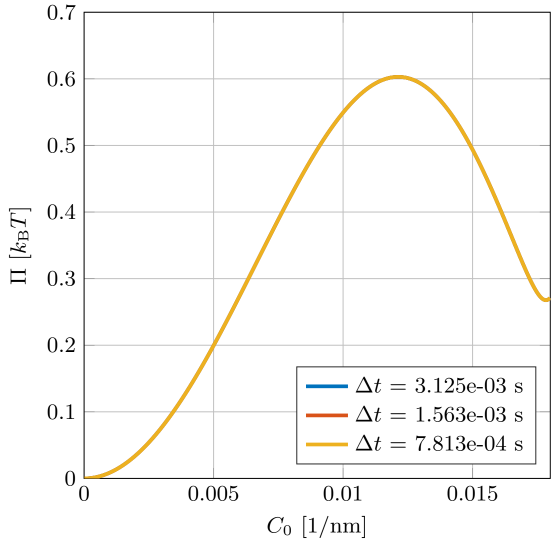

(a) at $\lambda_0 = 10^{-4}$ nm

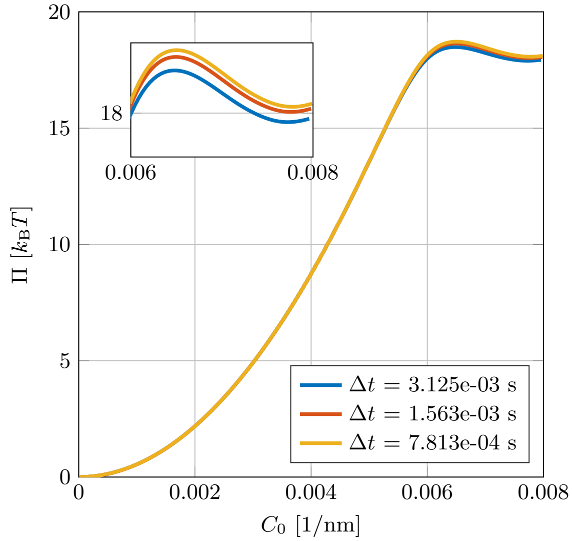

(b) at $\lambda_0 = 10^{-1}$ nm

Figure 11: Stored energy depending on the timestep size at different resting tensions in the non-axisymmetric setup.

##### 3.2.1 Elliptic vs. Circular Patches

In this section, we study the nature of the solutions arising from both circular and elliptic patches. A perfectly circular patch that induces curvature is not physically feasible, owing to thermal fluctuations or other inhomogeneities. Ellipticity is the first symmetry breaking perturbation of a circular region. It is for this reason that we use slightly elliptic patches of non-zero spontaneous curvature in studying the morphologies resulting from non-axisymmetric simulations.

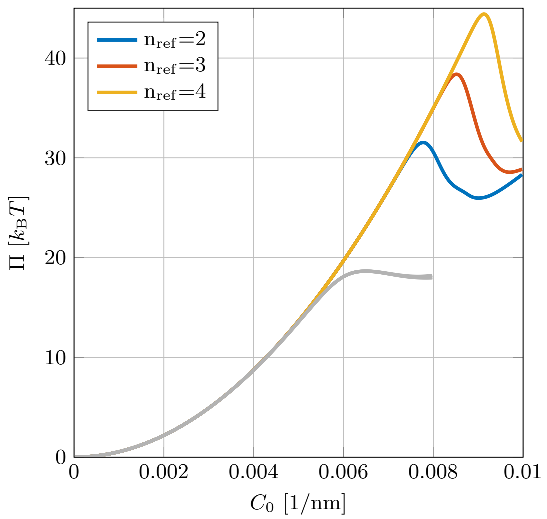

Figure 12: Convergence without an elliptic perturbation, at $\lambda_0 = 10^{-1}$ pN/nm. The gray line corresponds to the converged energy with an elliptic perturbation ($\delta = 0.02$).

Figure 12 shows the elastic energy (97) at different mesh refinement steps, when the coated area is circular. The peaks in energy indicate where the solution branches into non-axisymmetric shapes. Such branching is mesh-dependent as circular shapes can numerically only be approximated. Hence, initially circular patches are subject to mesh-dependent effects. Note that, with finer meshes, the numerical error associated with a circular patch decreases, and though instabilities still arise, branching is only observed at higher spontaneous curvatures $C_0$. Thus, we conclude that circular patches, due to their inherent symmetry, prolong the axisymmetric shapes leading to higher energies, and become unstable at sufficiently large spontaneous curvature $C_0$. Furthermore, the ridge morphology resulting from circular patches is shaped like a plus sign, such that the two lines are aligned with mesh lines (Fig. 13). This indicates that the ridge morphology is determined

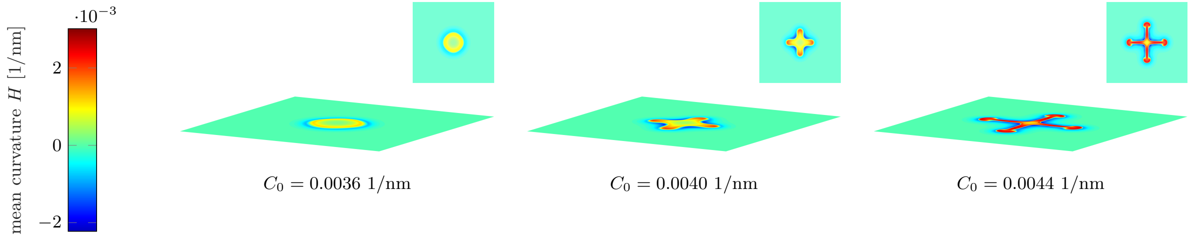

Figure 13: Plus shaped ridges resulting from an unperturbed circular coated region.

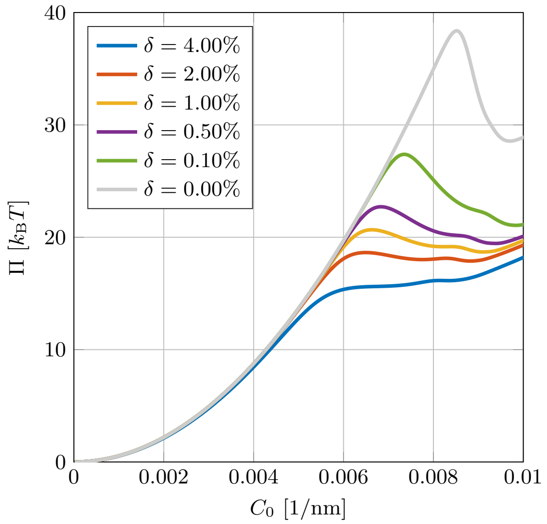

Figure 14: Stored energy depending on the ellipticity. The lengths of the principal semi-axes is given by $R_0(1 \pm \delta)$.

by our choice of the mesh. We note here that the results in which buds are forming are largely independent of the imposed ellipticity.

Finally, we comment on the magnitude of the ellipticity of the coated region, as slightly elliptical coated regions were used in the main text. The principal semi-axes of the coated region are given by

$$a = (1 + \delta)R_0 \ , \qquad b = (1 - \delta)R_0 \ , \tag{98}$$

and the elastic energy corresponding to different values of $\delta$ is plotted in Fig. 14. With increasing $\delta$, the branching point is not accompanied by a peak in the elastic energy. While all branches correspond to the formation of ridges, the exact values of the branching points are dependent on $\delta$: for small $\delta$, the pits remain axisymmetric to higher spontaneous curvatures—approaching the case of strictly circular patches. The latter preserve axisymmetry, leading to higher energies.

##### 3.2.2 Twisted Buds

We revisit the convergence behavior discussed in Fig. 8a at a low resting tension, now considering a larger range of the spontaneous curvature $C_0$. We find that the solutions do not converge after some value of $C_0$ as shown in Fig. 15. An example of the resulting shape is shown in Fig. 16, where we find the bud begins to twist about its vertical axis. When the spontaneous curvature is increased further, the twist of the neck becomes more pronounced until the buds shoot up into a vertical tube. Since such results are not converged, they require further careful study. Recall that in the high resting tension case, we introduced an elliptic perturbation to obtain stable shapes. Here, however, it is not clear what the required symmetry breaking perturbation is to obtain converged results at high spontaneous curvatures in the low resting tension case. Thus, this result is excluded from the main text.

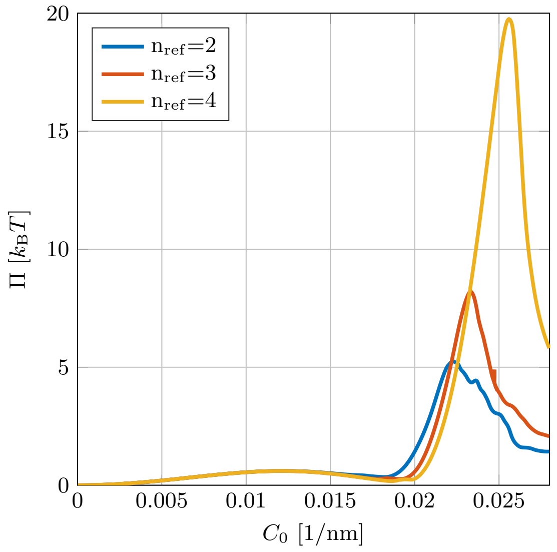

Figure 15: Stored energy depending the number of refinement steps in the center of the computational domain at $\lambda_0 = 10^{-4}$ nm, for non-axisymmetric solutions. The figure shows the stored energy beyond the values of spontaneous curvature where results are converged.

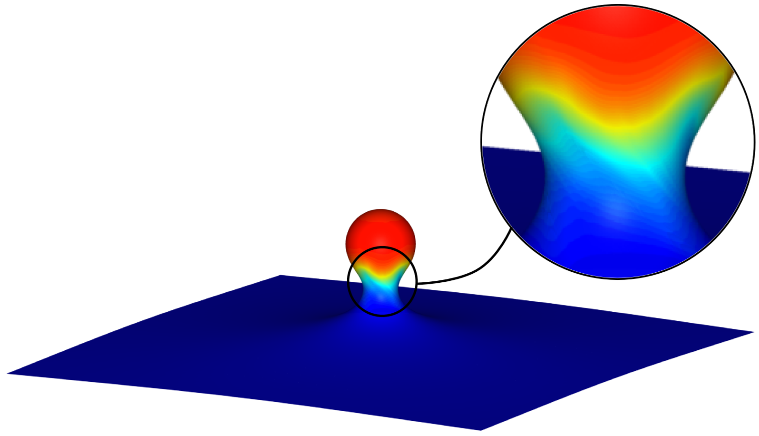

Figure 16: Close-up of the neck of a closed bud. The inset reveals that the spherical bud is twisted with respect to the flat lipid bilayer. Hence, it is apparent that the closed bud is not axisymmetric as well.

### 4 Additional results

#### 4.1 Fully Closed Axisymmetric Buds

In the main text, for the sake of comparison with the non-axisymmetric results, the axisymmetric results were shown at a spontaneous curvature $C_0$ for which the bud is not yet fully closed. In general, we find non-axisymmetric membrane morphologies in which the neck twists (see Fig. 16), while in axisymmetric simulations, the neck continues to constrict further (see Fig. 17). We find the bud is nearly spherical in shape, and the neck continues to tighten until the bud closes off—at which point our axisymmetric continuum simulations are no longer valid.

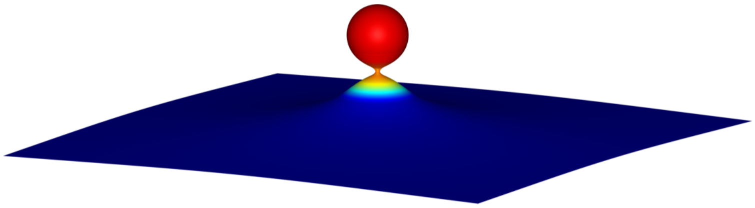

Figure 17: Closed bud in the axisymmetric setup at $\lambda = 10^{-4}$ pN/nm and $C = 0.023$ 1/nm. The neck is becoming strongly constricted, as opposed to the non-axisymmetric setup.

#### 4.2 Ridges at Lower Resting Tensions

In the main text, shallow ridges were reported at high resting tension. However, as the resting tension is lowered and approaches its threshold value $\lambda_0^{\text{thresh}}$, a taller ridge develops as shown in Fig. 18. The cylindrical nature of the ridge is more apparent in such cases.

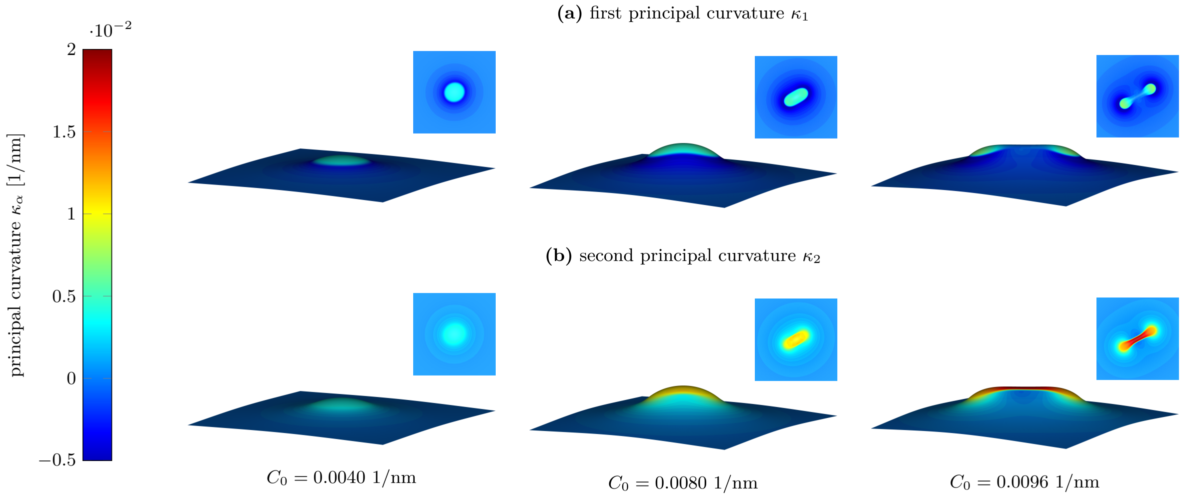

Figure 18: Plots of the principal curvatures of the non-axisymmetric shapes at $\lambda_0 = 3 \times 10^{-4}$ pN/nm. The chosen resting tension is just above the spontaneous curvature where ridges are observed, with $\dot{C}_0 = 0.1\dot{C}_0^0$. Here, we find that $H \approx C$ and thus $\kappa_2 \approx 2C$.
